# An Organoid for Woven Bone

**DOI:** 10.1101/2020.06.15.152959

**Authors:** Anat Akiva, Johanna Melke, Sana Ansari, Nalan Liv, Robin van der Meijden, Merijn van Erp, Feihu Zhao, Merula Stout, Wouter H. Nijhuis, Cilia de Heus, Claudia Muñiz Ortera, Job Fermie, Judith Klumperman, Keita ito, Nico Sommerdijk, Sandra Hofmann

**Author notes:** equal contribution (Anat Akiva, Johanna Melke).

## Abstract

Bone formation (osteogenesis) is a complex process in which cellular differentiation and the generation of a mineralized organic matrix are synchronized to produce a hybrid hierarchical architecture. To study the mechanisms of osteogenesis in health and disease, there is a great need for functional model systems that capture in parallel both cellular and matrix formation processes. Stem cell-based organoids are promising as functional, self-organizing 3D *in vitro* models for studying the physiology and pathology of various tissues. However, for human bone, no such functional model system is yet available.

This study reports the *in vitro* differentiation of human bone marrow stromal cells into a functional 3D self-organizing co-culture of osteoblasts and osteocytes, creating an organoid for early stage bone (woven bone) formation. It demonstrates the formation of an organoid where osteocytes are embedded within the collagen matrix that is produced by the osteoblasts and mineralized under biological control. Alike *in vivo* osteocytes the embedded osteocytes show network formation and communication via expression of sclerostin. The current system forms the most complete 3D living *in vitro* model system to investigate osteogenesis, both in physiological and pathological situations, as well as under influence of external triggers (mechanical stimulation, drug administration).

## 1. Introduction

Bone formation (osteogenesis) is a complex process in which i) cellular differentiation and ii) the generation of a mineralized organic matrix are synchronized to produce a hybrid hierarchical architecture.^[1]^ To study the molecular mechanisms of osteogenesis in health and disease there is great need for functional self-organizing 3D model systems that capture in parallel both cellular and matrix formation processes. Such a self-organizing 3D model system where mechanical and (bio)chemical signals can be applied in a dynamic environment, would be an important tool in the development of treatments for bone-related human diseases such as osteoporosis and osteogenesis imperfecta.

Organoids have been defined as “in vitro 3D cellular clusters derived exclusively from embryonic stem cells, induced pluripotent stem cells or primary tissue, capable of self-renewal and self-organization, and exhibiting similar organ functionality as the tissue of origin”, where they may “rely on artificial extracellular matrices (ECM) to facilitate their self-organization into structures that resemble native tissue architecture”.^[2]^ As the development of organoids relies on their self-organizing nature, they also often show variability in their results. Nevertheless, currently organoids are those *in vitro* model systems that most closely resemble the *in vivo* situation in tissues and provide a promising approach towards personalized medicine. However, for human bone, no such functional organoid is yet available.

A crucial challenge here is the realization of a 3D system with different interacting bone cell types. In particular the differentiation of *human* bone marrow-derived mesenchymal stromal cells (BMSCs) into osteocytes, which form 90-95% of the cellular fraction of bone tissue,^[3]^ has not yet been achieved *in vitro* and currently remains a critical step in the engineering of *in vitro* human bone models.

*In vivo*, osteocytes form through the differentiation of osteoblasts, after these become embedded in the extracellular matrix that they produce.^[3-4]^ Osteocytes are responsible for sensing the biophysical demands placed on the tissue and for orchestrating the concomitant actions of osteoblasts and osteoclasts in the remodeling of bone,^[1]^ as well as for maintaining calcium and phosphate homeostasis. During the differentiation from osteoblasts to osteocytes, the cells grow long extensions called processes, by which they form a sensory network that translates mechanical cues into biochemical signals and through which they interact with other cells.^[1]^

*In vitro*, osteoblast-based cell lines developed as models of osteocytes or osteocyte differentiation have not yet been shown to produce a fully developed mineralized collagen matrix,^[5]^ and hence are limited in their function as 3D models for bone formation. So far, the full differentiation from MSCs into functional osteocytes has been demonstrated for mouse cells^[6]^ but not yet for human cells. Recently, pre-osteocyte-like cells have been achieved from human primary cells,^[7]^ and co-cultures were generated from pre-prepared populations of osteoblasts and osteocytes,^[8]^ but the *in vitro* production of a bone-like mineralized matrix formed under biological control was not yet demonstrated. Also missing is a demonstration of the production of sclerostin by osteocytes at the protein level, where sclerostin is a key anti-anabolic molecule that interacts with osteoblasts to down-regulate ECM formation. Hence, the creation of an organoid, as a model for developing bone, through the full differentiation of human primary cells into a functional osteocyte network within a bone-like mineralized matrix is still an outstanding challenge.

As part of our efforts to realize a fully functional *in vitro* bone model, this work reports the differentiation of human BMSCs into a functional 3D self-organizing co-culture of osteoblasts and osteocytes, creating an organoid for early stage bone (woven bone) formation. We use a combination of immunohistochemistry, 2D and 3D electron microscopy and spectroscopy to demonstrate that the osteocytes form a network showing cell-cell communication via the expression of sclerostin, embedded within the collagen matrix that is formed by the osteoblasts and mineralized under biological control.

With this extensive characterization we demonstrate that this is the first fully functional 3D living *in vitro* model system for investigating the differentiation and matrix development processes during early bone formation.

## 2. Results

### 2.1. Differentiation of hBMSC into a 3D co-culture of osteoblast and osteocytes

In the present work, primary human bone marrow stromal cells (hBMSC) were seeded on porous 3D silk fibroin scaffolds^[9]^ and subsequently cultured in osteogenic differentiation medium. The cells were exposed to mechanical stimulation through fluid flow derived shear stress, by applying continuous stirring in a spinner-flask bioreactor (**Figure S1**), while a static system was used as control. Cells subjected to mechanical loading showed the production of a mineralized extracellular matrix (ECM) - as assessed by µCT and histological staining for collagen, glycosaminoglycans and minerals (**Figure S2**). FTIR spectroscopy indicated a matrix composition similar to that of embryonic chicken bones (**Figure S3**),^[10]^ In contrast, no significant ECM production was observed in the static system.

The histological sections also showed that the cells migrated within the scaffold, forming separated colonies. As these colonies present local environments, we expect the cells – as in any tissue - to show local variation in their degree of differentiation, related to differences in local mechanical stimulation, local cell density, scaffold shape or location within the scaffold.^[11]^ The simultaneous presence of several different developmental stages precludes monitoring cellular differentiation by standard genetic screening methods such as qPCR. The differentiation from primary cells to osteoblasts and osteocytes was therefore followed using immunohistochemistry, visualizing the expression of specific biomarkers at the protein level for the subsequent development to pre-osteoblasts, osteoblasts and osteocytes (**Figure 1, Figure S4**).^[1]^ The pre-osteoblastic stage was identified by the expression of transcription factors RUNX2 (CBFA-1) and osterix (OSX, SP7) (Figure 1 a-b, Figure S4). The next stage in the differentiation, the formation of osteoblasts, was heralded by the detection of osteoblast-specific markers, where alkaline phosphatase (ALP) was detected at the cell surfaces, and osteocalcin (BGLAP), osteopontin (BSP1) and osteonectin (SPARC) localized in the cellular environment (Figure 1 c-f, Figure S4). Finally, the differentiation into osteocytes was indicated by the expression of dentin matrix protein1 (DMP1), podoplanin (E11) and sclerostin (Figure 1 g-i, Figure S4). DMP1 is a marker for the early stages of osteocyte formation, coinciding with the embedding of the osteoblasts in the collagenous matrix. Podoplanin marks the osteocyte embedded in the non-mineralized matrix stage and has been suggested to regulate cell process formation.^[3]^ Sclerostin indicates the maturation of the osteocytes and their ability to perform their signaling function in the bone regulatory process.^[1, 12]^ We note that the rate of osteocyte differentiation depended not only on the exposure to mechanical stimulation, but also to the glucose concentration in the medium, as indicated by the detection of sclerostin after ∼4 weeks for 5.55 mM glucose (Figure 1 i) and ∼ 8 weeks for 25 mM glucose (Figure 1 k).

**Figure 1:**
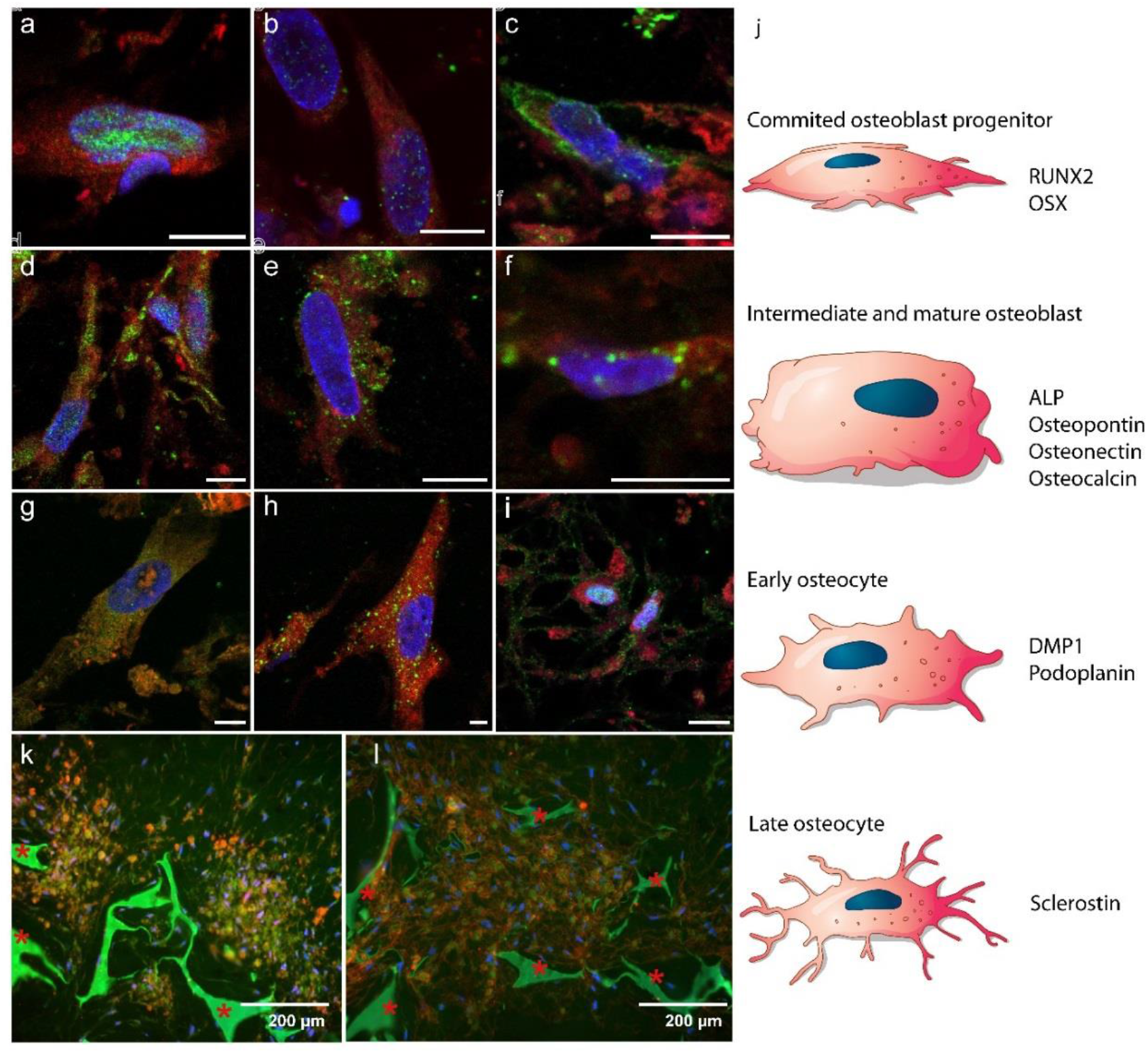
Differentiation of hBMSCs into osteoblasts and osteocytes: a-i) Fluorescence immunohistochemistry imaging showing markers for a-c) early stages of osteoblast formation, d-f) mature osteoblasts, and g-i) osteocyte development (5.6 mM glucose). Color code: red -cell cytoplasm, blue - cell nuclei, green: a) RUNX2 (day 7), b) OSX (day 7), c) ALP (day 26), d) osteocalcin (day 26), e) osteopontin (day 26), f) osteonectin (day 21), g) DMP1 (day 28), h) podoplanin (day 28), i) sclerostin (day 28). Scale bars: 10 µm. See Figure 4S for separate channels. j) Schematic illustration of MSC differentiation into osteoblasts and osteocytes, indicating at which state which protein expression is expected in a-i. k,l) Fluorescent images indicating self-organized domains of osteocytes embedded in a mineralized matrix after 8 weeks (25 mM glucose), k) co-localization of osteocytes (sclerostin, red) and mineral (calcein, green) and l) collagen (CNA35, red) and mineral (calcein, green) * Indicates the silk fibroin scaffold.

We then used fluorescence microscopy to investigate the ability of the cells to self-organize and to form a mineralized ECM. Staining of the mineral with calcein showed the co-localization with sclerostin in large domains with dimensions of hundreds of micrometers, indicating the embedding of the osteocytes in a mineralized ECM (Figure 1 k). Combined staining with calcein (mineral) and CNA35 (collagen) confirmed the presence of sub-millimeter sized mineralized collagen domains (Figure 1 l) in the pores of the scaffold throughout its entire volume. These mineralized domains co-existed with non-mineralized domains that stained positive for osteoblast markers (**Figure S5**).

This implies that a co-culture had formed in which osteogenic cells had organized themselves according to their stage of differentiation and maturation, and in which the osteocytes had become embedded within the mineralized matrix also produced by the system.

### 2.2 Osteocyte network analysis

Although the detection of DMP1, podoplanin and sclerostin markers indicated the presence of osteocytes in the co-culture, these observations do not prove that they form a connected functional network of cells. Fluorescence microscopy however confirmed the typical osteocyte morphology (**Figure 2**), showing the development of cell processes of > 10 micrometer, as well as the formation of an interconnected network (Figure 2 a, **Figure S6**). Additionally, 3D focused ion beam/scanning electron microscopy (3D FIB/SEM) showed that the cells form a relatively dense 3D network with significant variation in their morphologies, as well as in the number, length and connectivity of their processes (Figure 2 b-e, **Table S1 and S2**). We note that our osteocytes most often show flattened morphologies, which differ from those in text books with generally spherical or oblate bodies and long homogeneous protrusions, but are similar to osteocyte morphologies observed in different bone types, including rat tibia,^[13]^ human femur^[14]^ and mouse woven bone.^[15]^

**Figure 2:**
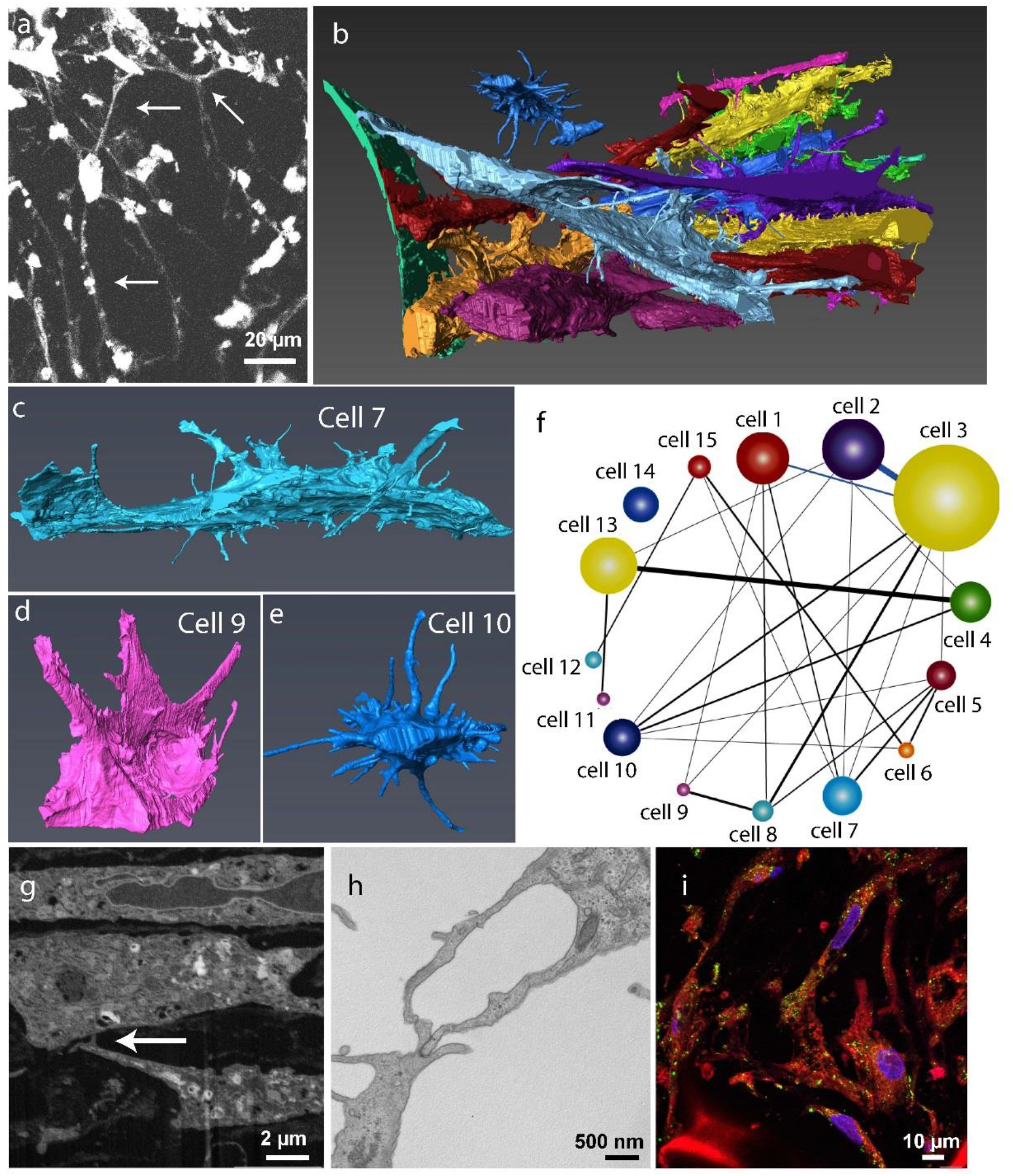
Osteocyte development and network formation. a) Fluorescent cytoplasm staining showing the development of long (> 5μm, arrows) cell processes connecting cells, and the formation of an interconnected network. The long processes were enhanced using a gamma value of -1.5. (see original image in Figure S6). b) 3D FIB/SEM reconstruction showing cell morphology and network formation in the whole volume. c-e) Details from the 3D reconstruction in (b) showing individual osteocytes in different stages of morphological development (cell numbers refer to identification in Figure S7). c) Cell #7, d) cell #9, e) cell #14. f-i) Cell connectivity: f) Connectivity map of the cells in the 3D FIB/SEM stack (see also Table S1 and S2). Sizes of the circles reflect the number of processes of that respective cell. Thickness of the lines reflects the number of connections between individual cells (cell numbers refer to identification in Figure S7). g) single slice from the 3D FIB/SEM stack showing long processes (arrow), creating a cellular network. h) TEM image shows a gap junction between processes of two connecting cells. i) fluorescent immunohistochemistry showing the presence of gap junctions on the surface of the different cells. Color code: red - cell cytoplasm, blue - cell nuclei, green - connexin43.

Cells showed both connected and unconnected processes, with connections to 1-7 neighboring cells (Figure 2 f, **Figure S7** and Table S2). The network had a density of 750,000 cells.mm^-3^, which is higher than observed for mature osteocytes in cortical bone (20,000-80,000 cells.mm^-3^),^[4]^ but in line with numbers found for woven bone such as in embryonic chicken tibia (500,000-700,000 cells.mm^-3^).^[16]^ Image analysis (**Figure S8**) showed that for the different cells the number of processes per unit surface area ranged between 0.05 - 0.23 µm^-2^ (**Table S3**), in line with values reported for mouse osteocytes (0.08 – 0.09 μm^-2^).^[17]^ The functionality of the processes was indicated not only by their connection to neighboring cells (Figure 2 g-h), but also by their co-localization with connexin43, a protein essential for gap junction communication (Figure 2 i).^[1]^ The observed variation in osteocyte morphology, together with the variation in the number of cell processes per surface area reflects the different stages of development and maturation, as expected in a differentiating osteogenic co-culture.

Hence, the osteocytes in our 3D in vitro system not only have the ability to organize themselves as a connective cellular network within a mineralized collagen matrix, but also show the capability to perform cell-cell communication.

### 2.3. Characterization of the extracellular matrix

Although it is essential that a 3D *in vitro* model for bone formation shows the relevant developmental stages, self-organization and cell-cell communication, its physiological relevance critically depends on its capability to reproduce the bone extracellular matrix. We therefore investigated the ability of our living *in vitro* system to form a functional extracellular matrix, in which collagen is mineralized under biological control (**Figure 3**). FIB/SEM volume imaging with 3D reconstruction showed that the osteocytes were fully embedded in their ECM (Figure 3 a; **Video S1**). The produced collagen matrix enveloped the cells, but showed a low degree of long range order (Figure 3 b), as known for woven bone.^[15, 18]^ and in line with what was described for collagen layers containing osteocytes.^[19]^ The extracellular deposition of non-collagenous proteins (NCPs) was evidenced by immunohistochemical analysis, showing the presence of osteocalcin, osteopontin, and DMP1 and their co-localization with collagen (Figure 3 c-d, **Figure S9**).

**Figure 3:**
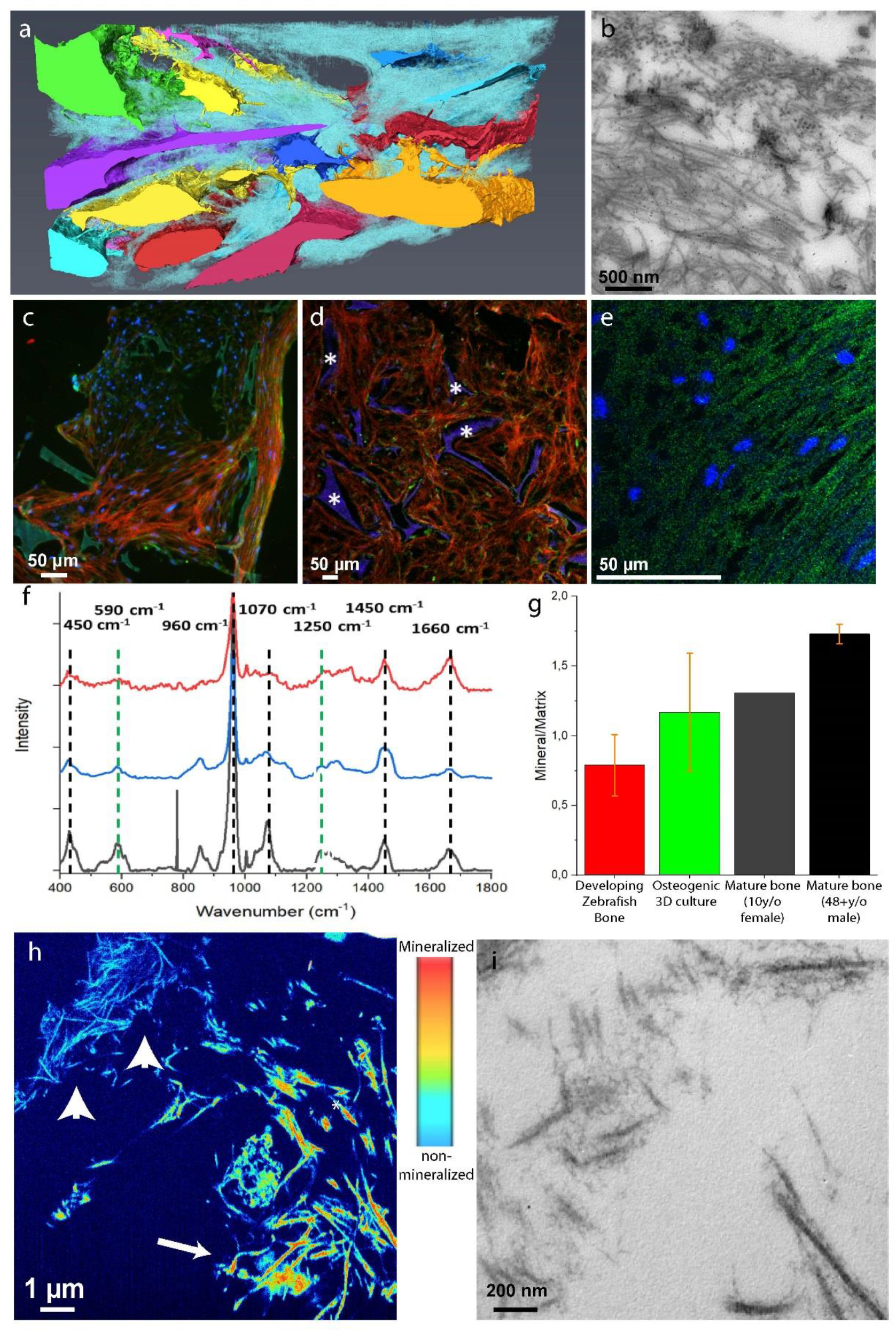
Extracellular matrix development: a) 3D FIB/SEM reconstruction shows the embedding of the cells in the collagen matrix (cyan). Discrete cells are represented with different colors. b) TEM image of a 70 nm section showing the random distribution of collagen fibrils. Collagen type I was identified by immunolabelling. c-e) Fluorescent immunohistochemistry identifying key non-collagenous proteins in the collagenous matrix: c) Co-localization of osteocalcin (green) and collagen (red). d) Osteopontin (green) distribution in the collagen matrix (red). * Indicates the silk fibroin scaffold. e) Co-localization of DMP1 (green) with the collagen structure (see Figure S5 for collagen image). f-g) Raman microspectrometry of mineralized matrices. f) Localized Raman spectra of mineralized collagen of developing zebrafish bone (red), the 3D osteogenic co-culture (blue) and human bone of a 10 year old female (grey g) Raman derived mineral/matrix ratios of 4 mineralized tissues of zebrafish (N=6, red), Osteogenic 3D culture (N=7, green), 10 year old human female (N=1, grey), and 48+ years old human male (N=7, black, taken from reference^[33]^). Bars indicate sample standard deviations. h) Heat map presentation of a 3D FIB/SEM cross section showing collagen fibrils with different degrees of mineralization (Figure S10). Arrowheads indicate non-mineralized collagen fibrils (light blue), arrow indicates mineralized collagen fibril (orange) i) TEM image showing individual mineralized collagen fibrils.

Whereas µCT (Figure S2), FTIR (Figure S3), histochemistry (Figure S2), and fluorescence microscopy (Figure 1 k) all indicated the mineralization of the organic matrix, none of these methods can provide the spatial resolution to demonstrate whether the mineral crystals are indeed, as in bone, co-assembled with the collagen fibrils,^[20]^ and not just the result of uncontrolled precipitation.^[21]^ We therefore applied a multiscale imaging approach to verify that matrix mineralization indeed occurred under biological control. Raman micro-spectroscopy of the extracellular matrix showed the spectral signature of developing bone (Figure 3 f) and confirmed the co-localization of the mineral with the collagen (**Figure S10**. Spectral analysis further confirmed that the mineral/matrix ratio (a key parameter for bone development, determined from the PO_4_ ν_4_ / Amide III vibrations intensity ratio ^[22]^) in the co-culture was indeed in the range found for developing bone (Figure 3 g).^[23]^

At higher resolution (voxel size 10×10×20 nm^3^), 3D FIB/SEM with back scatter detection revealed thin collagen fibrils (diameters 50-80 nm) with varying degrees of mineralization as also commonly observed in the early stages of bone development (Figure 3 i, **Video S2, Figure S11**).^[24]^ Applying a heat map presentation showed the coexistence of non-mineralized fibrils (blue) alongside a mineralized population (green-red range). Additionally, transmission electron microscopy (TEM) showed mineralized single fibrils,^[6a]^ indicating that the collagen matrix was indeed mineralized under biological control (Figure 3 j).^[25]^ Nevertheless, in some areas also larger mineral precipitates were observed (Figure S10 c, orange) (Video S2), possibly due to local non-biologically controlled precipitation of calcium phosphate.

## 3. Discussion

Woven bone is the first form of mammalian bone deposited during embryonic development and fracture, before being replaced by other bone types.^[18]^ Hence, in situations in which rapid formation is a prime concern, and where osteoclasts and bone remodeling do not yet play a role. Our results show the formation of a bone organoid consisting of a self-organized co-culture of osteoblasts and osteocytes representing a functional model for woven bone, an early state of bone formation in which the collagen matrix is still disorganized and distinct from the more mature ordered 3D structure.^[20]^ We demonstrate the functionality of the organoid by showing that the ECM formed by the osteoblasts is mineralized under biological control, and that the mature osteocytes self-organize into a network within the mineralized matrix where they express sclerostin and connexin43 at the protein level.

Interestingly, the production of sclerostin did not prohibit ECM formation throughout the organoid, suggesting that the down regulation of this process is a local effect. This may be explained by assuming the most mature osteocytes are in the center of the osteocyte domains, which would lead to a gradient of sclerostin decreasing towards the periphery of the network, only affecting the activity of the osteoblasts closest to the osteocyte domain.

The use of silk fibroin as a scaffold material rather than the frequently used collagen scaffolds, permits to differentiate between the supplied and the newly formed matrix material, and study the quality of the collagen matrix as function of external stimuli (mechanical load, therapeutics) or genetic diseases (e.g. osteogenesis imperfecta). The application of mechanical stimulation during the development of our stem cell based co-culture proved crucial for the osteogenic differentiation, and underlines the importance of the integration of self-organizing stem cell based strategies with environmental control in microfluidic systems in the recent organoid-on-a-chip approaches.^[26]^

## 4. Conclusion

Summarizing we conclude that we have generated an organoid for woven bone. The ability of this self-organizing 3D co-culture of osteoblasts and osteocytes to form an organic matrix that is mineralized under biological control is currently the most complete human *in vitro* model system for bone formation. It introduces the ability to closely monitor both cellular and matrix formation processes and will thereby provide new unique possibilities for the study of genetic bone related diseases, and the development of personalized medicine.

## 5. Experimental Section/Methods

### 5.1. Materials

Dulbecco’s modified Eagle medium (DMEM high glucose Cat. No. 41966 and low glucose Cat. No. 31885) and antibiotic/antimycotic (Anti-Anti) were from Life Technologies (Bleiswijk, The Netherlands). Citrate buffer was from Thermo Fisher Scientific (Breda, The Netherlands). Methanol was from Merck (Schiphol-Rijk, The Netherlands). Trypsin-EDTA (0.25%) was from Lonza (Breda, The Netherlands). Fetal bovine serum (FBS) was from PAA Laboratories (Cat. No A15-151, Cölbe, Germany). 10-nm Au particles conjugated to Protein-A were from CMC, UMC Utrecht (Utrecht, The Netherlands). BSA-c was from Aurion (Wageningen, The Netherlands). Silkworm cocoons from Bombyx mori L. were purchased from Tajima Shoji Co., LTD. (Yokohama, Japan). All other substances were of analytical or pharmaceutical grade and obtained from Sigma Aldrich (Zwijndrecht, The Netherlands). The reference human bone sample used for Raman micro-spectrometry was waste material from a surgical procedure on a fractured left tibia of a 10-year-old female. According to the Central Committee on Research involving Human Subjects (CCMO), this type of study does not require approval from an ethics committee in the Netherlands (see https://english.ccmo.nl/investigators/legal-framework-for-medical-scientific-research/your-research-is-it-subject-to-the-wmo-or-not). For information on the embryonic chicken bone and zebrafish bone we refer to references [15] (Kerschnitzky et al.) and [23] (Akiva et al.), respectively.

### 5.2. Scaffold fabrication

Silk fibroin scaffolds were produced as previously described^[27]^. Briefly, Bombyx mori L. silkworm cocoons were degummed by boiling in 0.2 M Na_2_CO_3_ twice for 1 h. The dried silk was dissolved in 9 M LiBr and dialyzed against ultra-pure water (UPW) for 36 h using SnakeSkin Dialysis Tubing (molecular weight cutoff: 3.5 K; Thermo Fisher Scientific, Breda, The Netherlands). Dialyzed silk fibroin solution was frozen at -80°C and lyophilized (Freezone 2.5, Labconco, Kansas City, MO, USA) for 4 days, then dissolved in hexafluoro-2-propanol, resulting in a 17% (w/v) solution. Dissolved silk fibroin (1 ml) was added to NaCl (2.5 g) with a granule diameter of 250–300 μm and was allowed to air dry for 3 days. Silk-salt blocks were immersed in 90% MeOH for 30 min to induce β-sheet formation^[28]^. NaCl was extracted from dried blocks in UPW for 2 days. Scaffolds were cut into disks of 5 mm in diameter and 3 mm in height and autoclaved in PBS at 121^°^C for 20 min.

### 5.3. Cell culture

Cells were isolated from unprocessed, fresh, human bone marrow (Lonza, Walkersville, MD, USA, cat. No #1M-125) of one male donor (healthy, non-smoker). hBMSC isolation and characterization was performed as previously described and passaged up to passage 4^[27]^. Pre-wetted scaffolds were seeded with 1 million cells each in 20 μL control medium (DMEM, 10% FBS, 1% Anti-Anti) and incubated for 90 min at 37°C. The cell-loaded scaffolds were transferred to custom-made spinner flask bioreactors (*n* = 4 per bioreactor, Figure S1). Each bioreactor contained a magnetic stir bar and was placed on a magnetic stirrer (RTv5, IKA, Germany) in an incubator (37°C, 5% CO_2_). Each bioreactor was filled with 5 mL osteogenic medium (control medium, 50 μg.mL^-1^ ascorbic-acid-2-phosphate, 100 nM dexamethasone, 10 mM β-glycerophosphate) and medium was changed 3 times a week.

### 5.4. Micro-computed tomography imaging (µCT)

μCT measurements and analysis were performed on a μCT100 imaging system (Scanco Medical, Brüttisellen, Switzerland). Scanning of the co-culture samples within the bioreactor was performed at an isotropic nominal resolution of 17.2 μm, energy level was set to 45 kVp, intensity to 200 μA, 300 ms integration time and two-fold frame averaging. A constrained Gaussian filter was applied to reduce part of the noise. Filter support was set to 1.0 and filter width sigma to 0.8 voxel. Filtered grayscale images were segmented at a global threshold of 23% of the maximal grayscale value to separate the mineralized tissue from the background and binarize the image. Unconnected objects smaller than 50 voxels were removed and neglected for further analysis. Quantitative morphometrical analysis was performed to assess mineralized ECM volume within the entire scaffold volume using direct microstructural bone analysis as previously described for human bone biopsies^[29]^.

### 5.5. Histological analysis

#### 5.5.1. Fixation and sectioning

Co-cultures were fixed in 10% neutral buffered formalin (24 h at 4° C), dehydrated in serial ethanol solutions (50%, 70%, 90%, 96%, 100%, 100%, 100%), embedded in paraffin, cut into 6 μm thick sections and mounted on Poly-L-Lysine coated microscope slides. Paraffin sections were dewaxed with xylene and rehydrated to water through graded ethanol solutions.

#### 5.5.2. Brightfield imaging

Sections were stained with Alizarin Red to identify mineralization (2%, Sigma-Aldrich), Picrosirius Red (0.1%, Sigma-Aldrich) to identify collagen, Alcian blue (1%, Sigma-Aldrich) to identify Glycosaminoglycan (GAGs). Sections were imaged using Zeiss Axio Observer Z1 microscope.

#### 5.5.3. Immunohistochemistry

Antigen retrieval in pH 6 citrate buffer at 95° C was performed for 20 min. Sections were washed three times in PBS. Non-specific antibody binding was blocked with 5% serum (v/v) from the host of the secondary AB and 1% bovine serum albumin (w/v) in PBS (blocking buffer) for 1 h. Sections were then incubated overnight at 4° C with primary antibodies in blocking buffer. The sections were rinsed with PBS four times for 5 min and incubated for 1 h with secondary antibodies in blocking buffer and at times with calcein solution (1 μg.mL^-1^, C0875 Sigma-Aldrich). All used antibodies and dyes are listed in Table 1. Nuclei were stained with DAPI for 5 min, after which sections were again washed three times with PBS and mounted on microscope glass slides with Mowiol. Cytoplasm was stained with FM 4-64 (Molecular Probes cat#T3166) for 1 minute and followed by washing the sections three times with PBS. Except for primary antibody incubation, all incubation steps were performed at room temperature. Sections were imaged either by Zeiss Axiovert 200M microscope (large field of view) or by Leica TCS SP5X (x63, high magnification images). Images were post processed (brightness, contrast, channel merging and crop) using Fiji software.

**Table 1.** List of all antibodies and dyes used.

| **Antigen** | **Source** | **Cat. No** | **Label** | **Species** | **Dilution/concentration** |
| --- | --- | --- | --- | --- | --- |
| ALP | ThermoFisher | MA5-17030 | - | Mouse | 1:200 |
| Calcein Green | Sigma | C0875 | - | - | 1 $\mu\text{g.mL}^{-1}$ |
| CNA35-mCherry | Homemade<sup>[34]</sup> | - | mCherry | - | 1 $\mu\text{M}$ |
| Collagen1 | Abcam | ab34710 | - | Rabbit | 1:250 |
| Connexin 43 | Sigma | C6219 | - | Rabbit | 1:500 |
| DAPI | Sigma | D9542 | - | - | 0.1 $\mu\text{g.mL}^{-1}$ |
| DMP1 | biorbyt | orb247330 | - | Rabbit | 1:400 |
| FM 4-64 | Molecular Probes | T3166 | - | - | 5 $\mu\text{g.mL}^{-1}$ |
| Osteocalcin | Abcam | Ab93876 | - | Rabbit | 1:200 |
| Osteonectin | ThermoFisher | MA1-43027 | - | Mouse | 1:500 |
| Osteopontin | ThermoFisher | 14-9096-82 | - | Mouse | 1:200 |
| OSX | Abcam | ab22552 | - | Rabbit | 1:200 |
| Podoplanin | Abcam | ab128994 | - | Rabbit | 1:200 |
| RUNX2 | Abcam | Ab23981 | - | Rabbit | 1:500 |
| Sclerostin | ThermoFisher | PA5-37943 | - | Goat | 1:200 |
| Goat IgG (H+L) | Jackson Immuno | A11055 | Alexa 488 | Donkey | 1:200 |
| Mouse IgG (H+L) | Molecular Probes | A21236 | Alexa 647 | Goat | 1:200 |
| Mouse IgG (H+L) | Jackson Immuno | 715-545-150 | Alexa 488 | Donkey | 1:200 |
| Rabbit IgG (H+L) | Invitrogen | A21206 | Alexa 488 | Donkey | 1:200 |
| Rabbit IgG (H+L) | Molecular Probes | A21244 | Alexa 647 | Goat | 1:200 |
| Alizarin Red | Sigma-Aldrich | A5533 | - | - | 2% |
| Picrosirius Red | Sigma-Aldrich | 365548 | - | - | 0.1% |
| Alcian blue | Sigma-Aldrich | A5268 | - | - | 1% |

### 5.6. Electron microscopy

#### 5.6.1. Sample preparation for electron microscopy

Samples were processed for electron microscopy as previously described.^[30]^ In short, co-culture samples were fixed in 2.5% glutaraldehyde and 2% paraformaldehyde in 0.1 M sodium cacodylate buffer (CB) for 72 h and washed 5 times in 0.1 M CB and 5 times in double-distilled water (ddH_2_O). Co-cultures were then post fixed using 1% OsO_4_ with 0.8% K_3_Fe(CN)_6_ in 0.1 M CB for 1 hour on ice. After rinsing in 0.1 M CB, the co-cultures were treated with 1% Tannic acid followed by 1% uranyl acetate in ddH_2_O for 1 hour. Finally, the samples were rinsed using ddH_2_O, dehydrated with ethanol (50%, 70%, 90%, 96%, 100%), and embedded in Epon resin.

#### 5.6.2. Focused Ion Beam Scanning Electron microscopy (FIB/SEM) imaging

Epon embedded samples were imaged with a Scios FIB/SEM (Thermo Fisher Scientific, Breda, The Netherlands) under high vacuum conditions. Using the gas injection system (GIS) in the FIB/SEM microscope, a 500 nm thick layer of Pt was deposited over the ROI, at an acceleration voltage of 30 kV and a current of 1 nA. Trenches flanking the ROI were milled at an acceleration voltage of 30 kV, using a high FIB beam current (5-7 nA), followed by a staircase pattern in front of the ROI to expose the imaging surface. Fine polishing was performed with the ion beam set to 30 kV with a FIB beam current of 0.5 nA, resulting in a smooth imaging surface. Serial imaging was then performed using the in-column backscattered electron detector, and the following settings: Acceleration voltage 2 kV, Beam current 0.2 nA, Pixel dwell time 10 μs, voxel size: 30×30×30 nm (stack in video S1) and 10×10×20 nm (stack in Video S2).

#### 5.6.3. Sample preparation for Transmission Electron Microscopy (TEM)

Epon embedded samples: 70 nm sections from resin embedded blocks were made using an ultra-microtome (Leica), and collected on carbon coated copper TEM grids. Post staining with uranyl acetate and led citrate was performed using the Leica EM AC20 automatic contrasting instrument.

#### 5.6.4. Preparation and immunogold labelling of Tokuyasu Sections

Thin sections were prepared following the Tokuyasu protocol.^[31]^ Briefly, co-cultures were fixed as described above and infiltrated overnight in 2.3 M sucrose for cryo-protection. Small blocks of the co-cultures were mounted on aluminum pins and plunge frozen in liquid nitrogen. 70 nm thick cryosections were sectioned with a cryo-ultramicrotome and picked up with a mixture of 2% methylcellulose/2.3 M sucrose on copper support grids coated with formvar and carbon. After rinsing away the pick-up solution in PBS at 37° C for 30 minutes, the sections were treated with PBS containing 0.15% glycine, followed by blocking for 10 minutes with 0.5% cold fish skin gelatin and 0.1% BSA-c in PBS. The TEM grids were incubated for 1 hour with a collagen type 1 antibody in blocking solution (Abcam, AB34710). The grids were then rinsed with 0.1% BSA in PBS and incubated with 10-nm Au particles conjugated to Protein-A in blocking solution.^[32]^ The sections were then thoroughly washed in ddH_2_O, stained with uranyl acetate and embedded in methylcellulose.^[32]^

#### 5.6.5. TEM imaging

The sections were imaged using a Tecnai T12 TEM (80kV) (Thermo Fisher Scientific, Breda, The Netherlands), equipped with Veleta (EMSIS GmbH, Münster, Germany).

### 5.7. Raman Spectroscopy

Raman measurements were conducted using a WiTec Alpha 300R confocal Raman microscope. Co-culture samples were fixed in 10% neutral buffered formalin (24 h at 4° C), incubated for 2 hours in 5% sucrose at 4° C, embedded in Tissue-Tek (Sakura Finetek 4609024), cut into 10 μm thick sections and mounted on Poly-L-Lysine coated microscope slides. Raman imaging was conducted using 532 nm excitation lasers with a laser power of 10 mW, using 50x 0.8 objective (0.8 NA) with a grating of 600 mm^-1^. The maps were obtained with a spatial resolution of 3 spectra.µm^-1^. Data analysis was performed using Project V plus software (Witec, Ulm) and Origin 8.

### 5.8. FTIR Spectroscopy

Prior to the FTIR measurement, the co-culture samples were freeze dried overnight. After drying, 1.5 mg of the samples was crushed using mortar and pestle until a fine powder was achieved. After this, 148.5 mg of KBr was added to the mortar and pestle and the materials were mixed and further crushed to a fine homogeneous mixture. The mixture was added to a pellet press holder, the transparent and homogeneous pellets were then inserted into the FTIR spectrometer (Perking Elmer one 1600). The FTIR spectra were obtained in transmission mode. Spectra were obtained over the range form 200 cm^-1^ to 6000 cm^-1^ with a spectral resolution of 0.5 cm^-1^.

### 5.9. Image Analysis

3D FIB/SEM Image processing was performed using Matlab and Avizo 3D software (FEI VSG, www.avizo3d.com). 3D image reconstruction, alignment, denoising and brightness and contrast adjustments were done usign Matlab. 3D segmentation was done using Avizo. Segmentation was performed using manual thresholding, and cell processes were counted manually. As all cells were only partially captured in the available FIB/SEM volume, cell process density was determined per unit surface area of the cell body. The surface areas of cell parts captured in the FIB/SEM volume were calculated from the segmented 3D image mask using Matlab and Fiji. Further details are given in Fig. S11.

Cell density was determined from the number of cells in the imaged FIB/SEM volume (20 µm x 20 µm x 40 µm) and compared to literature data from histological sections (volume 5 µm x 1000 µm x 1000 µm).^[16]^

## Acknowledgements

We would like to thank like Lia Addadi and Steve Weiner for providing the zebrafish and embryonic chicken data, and Carlijn Bouten for providing CNA35 collagen probe. We also thank Deniz Daviran for her help in preparing the figures. AA was supported by the Marie Curie Individual Fellowship (H2020-MSCA-IF-2017-794296-SUPERMIN), by the Netherlands Organization for Scientific Research (NWO) through an ECHO grant to NS, and by the National Postdoctoral Award Program for Advancing Women in Science – the Weizmann Institute of Science, Israel. NS, RvdM and MvE were supported by the European Research Council (ERC) Advanced Investigator grant (H2020-ERC-2017-ADV-788982-COLMIN) to NS, SA was supported by the Ministry of Education, Culture and Science (Gravitation Program 024.003.013). NL was supported by the Netherlands Organization for Scientific Research (NWO) through a ZonMW-TOP grant to JK. JM, FZ and SH were supported by the ERC Starting grant (FP7-ERC-2013-StG-336043-REMOTE) to SH. AA and JM contributed equally to this work.

## Supporting Information

**Figure S1:**
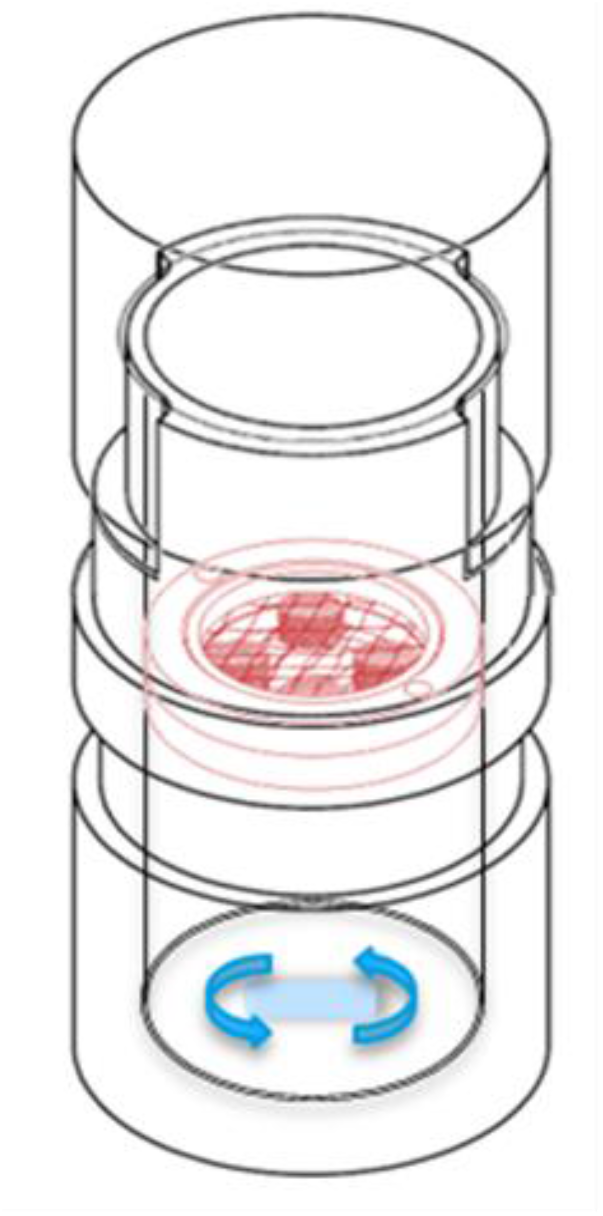
Schematic of the spinner flask type bioreactor adapted for longitudinal microCT scanning. The 3D silk fibroin scaffolds are seeded with the hBMSCs and fixed between two meshes in the inlay of the bioreactor (red). The constructs are continuously exposed to shear stress (see reference [11]) provided by a magnetic stirrer bar, located at the bottom of the bioreactor (blue). Dimensions of the bioreactor: outer diameter: 36mm, height: 80mm.

**Figure S2:**
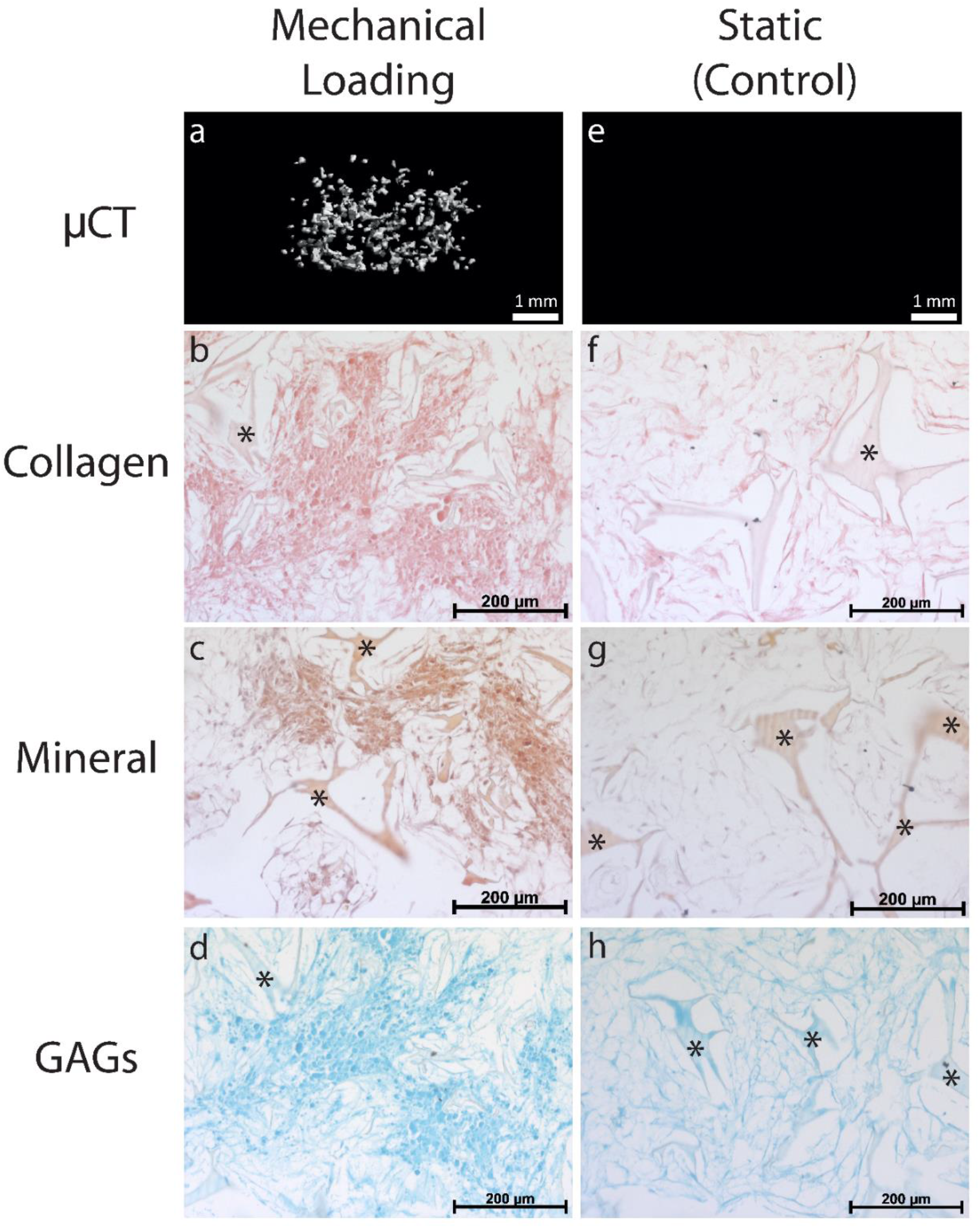
Characterization of the extracellular matrix produced by the cells in the construct (a-d) in a spinner flask bioreactor under mechanical loading and (e-h) in a static system. µCT image of deposited mineral (a,e), and histological identification of collagen (Picrosirius Red) (b,f), mineral (Alizarin Red)(c,g) and glycosaminoglycans (GAGs, Alcian blue)(d,h). The results show that mechanical loading through shear stress is critical for the development of the extracellular matrix. * denotes the silk fibroin scaffold.

**Figure S3:**
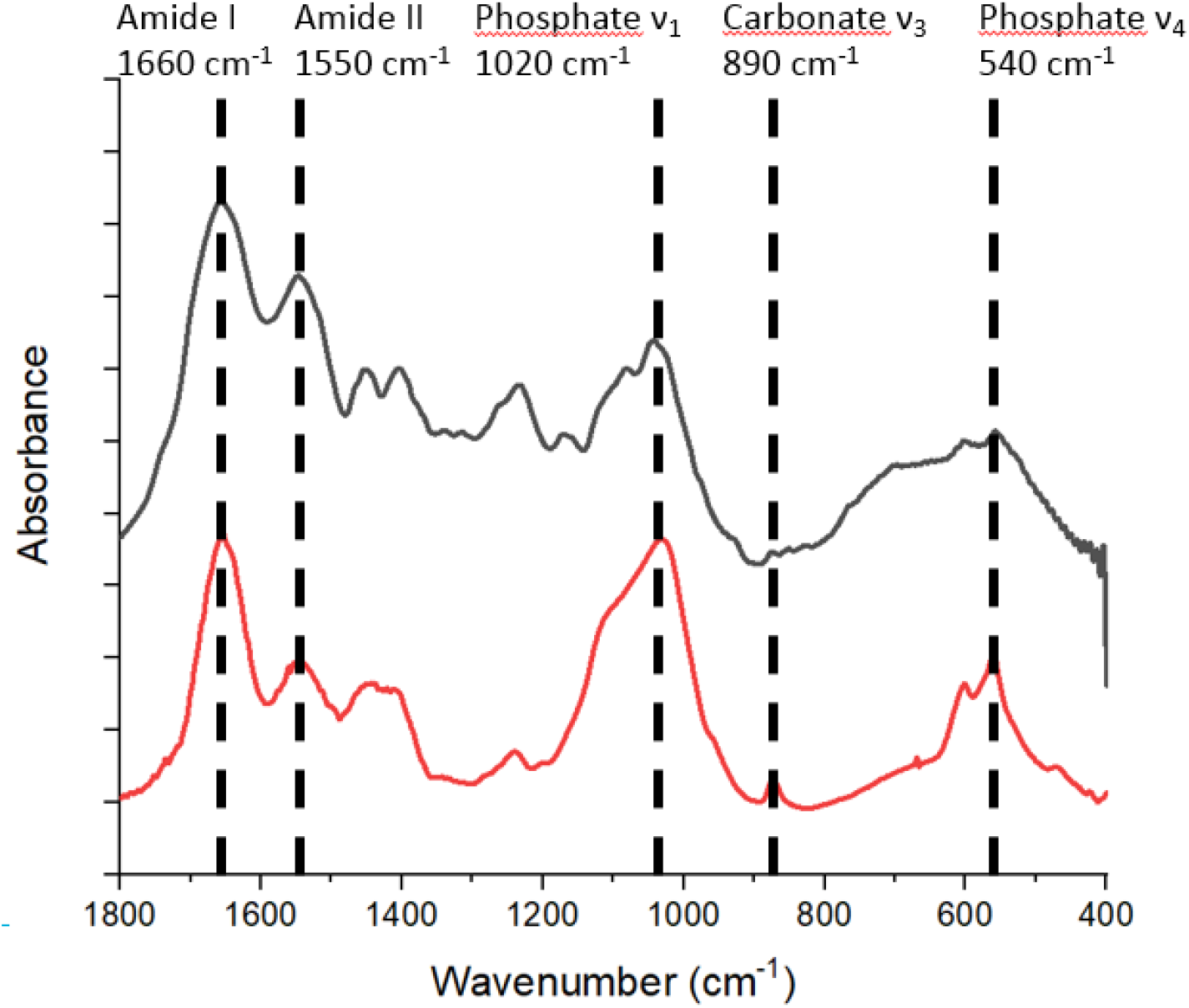
FTIR spectrum of the osteogenic 3D co-culture after 10 weeks of development (black line), and of embryonic young chicken long bones at embryonic stage E16(red line); see also reference [15] (Kerschnitzki et al.). The spectra show similar peak positions for the main organic and inorganic contributions indicated by the dashed lines, as well as similar Amide I /Phosphate ν_1_ peak intensities, which indicates both samples have similar levels of tissue mineralization. Spectra were baseline corrected via straight line fitting to the 400 cm^-1^ position, 1800 cm^-1^ and the minimum at 900 cm^-1^. The 3D osteogenic co-culture sample was corrected for the silk fibroin scaffold contributions approximated to be 40 wt%. When subtracting higher values for the scaffold contribution, negative peaks became evident in the spectra. Both spectra were normalized to the height of the Amide I band (1660 cm^-1^).

**Figure S4a:**
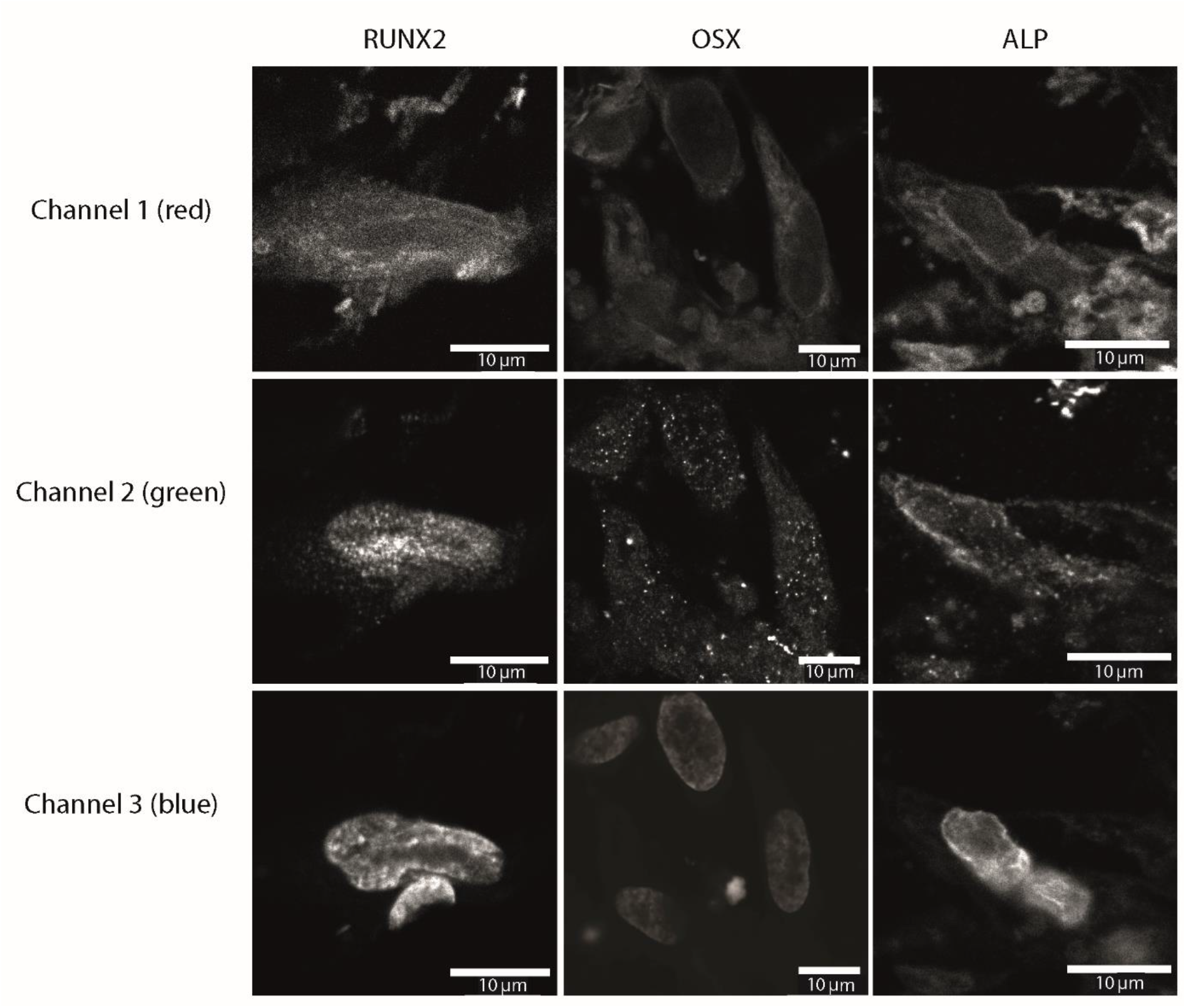
Separate channels of all panels from Figure 1. The information of the three individual channels (Channel 1 - red, cell cytoplasm, Channel 2 - green, targeted protein and Channel 3 -blue, nuclei) are shown separately in gray scale for each image in Figure 1 a-c.

**Figure S4b:**
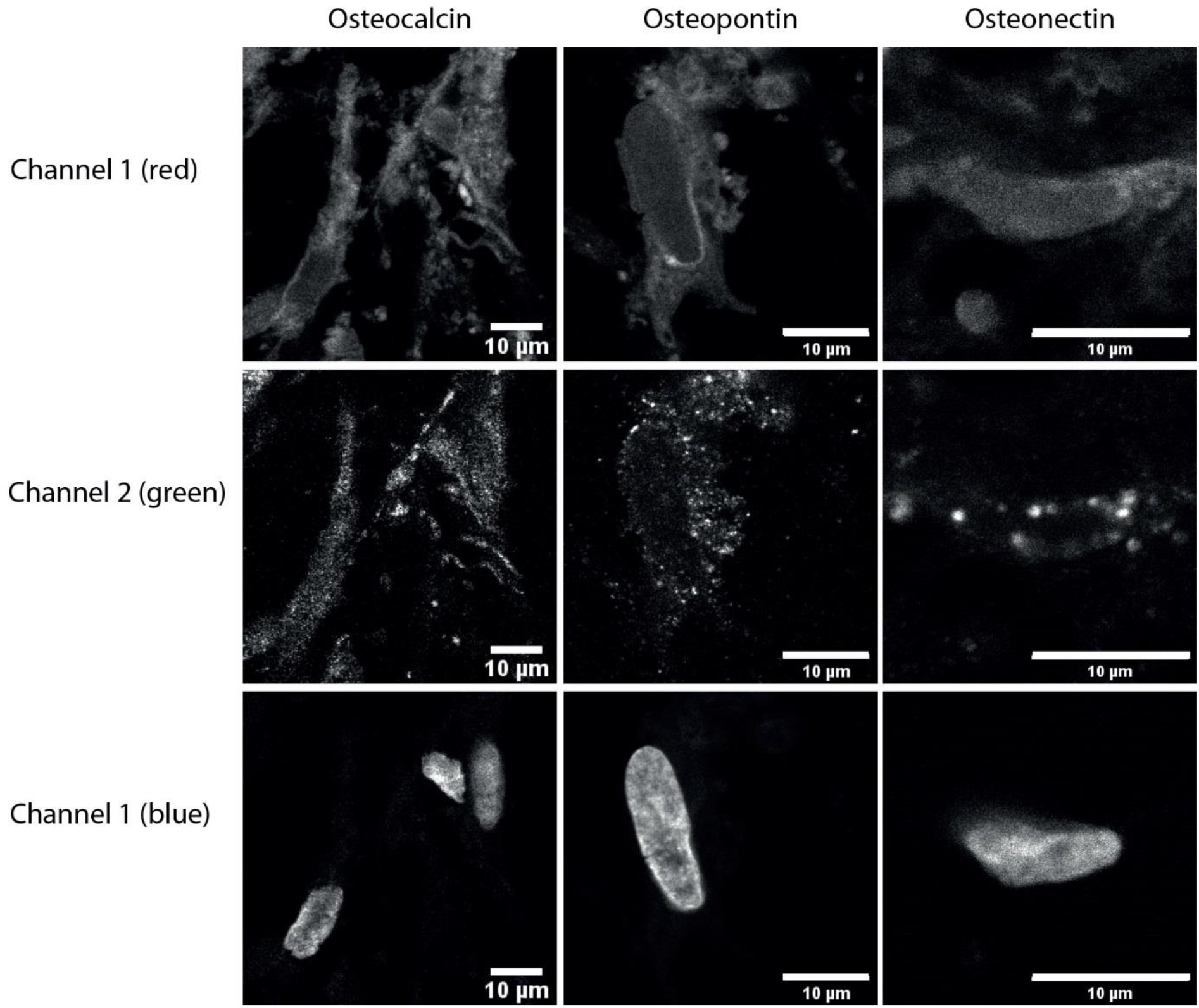
Separate channels of all panels from figure 1. The information of the three individual channels (Channel 1 - red, cell cytoplasm, Channel 2 - green, targeted protein and Channel 3 -blue, nuclei) are shown separately in gray scale for each image in Figure 1 d-f.

**Figure S4c:**
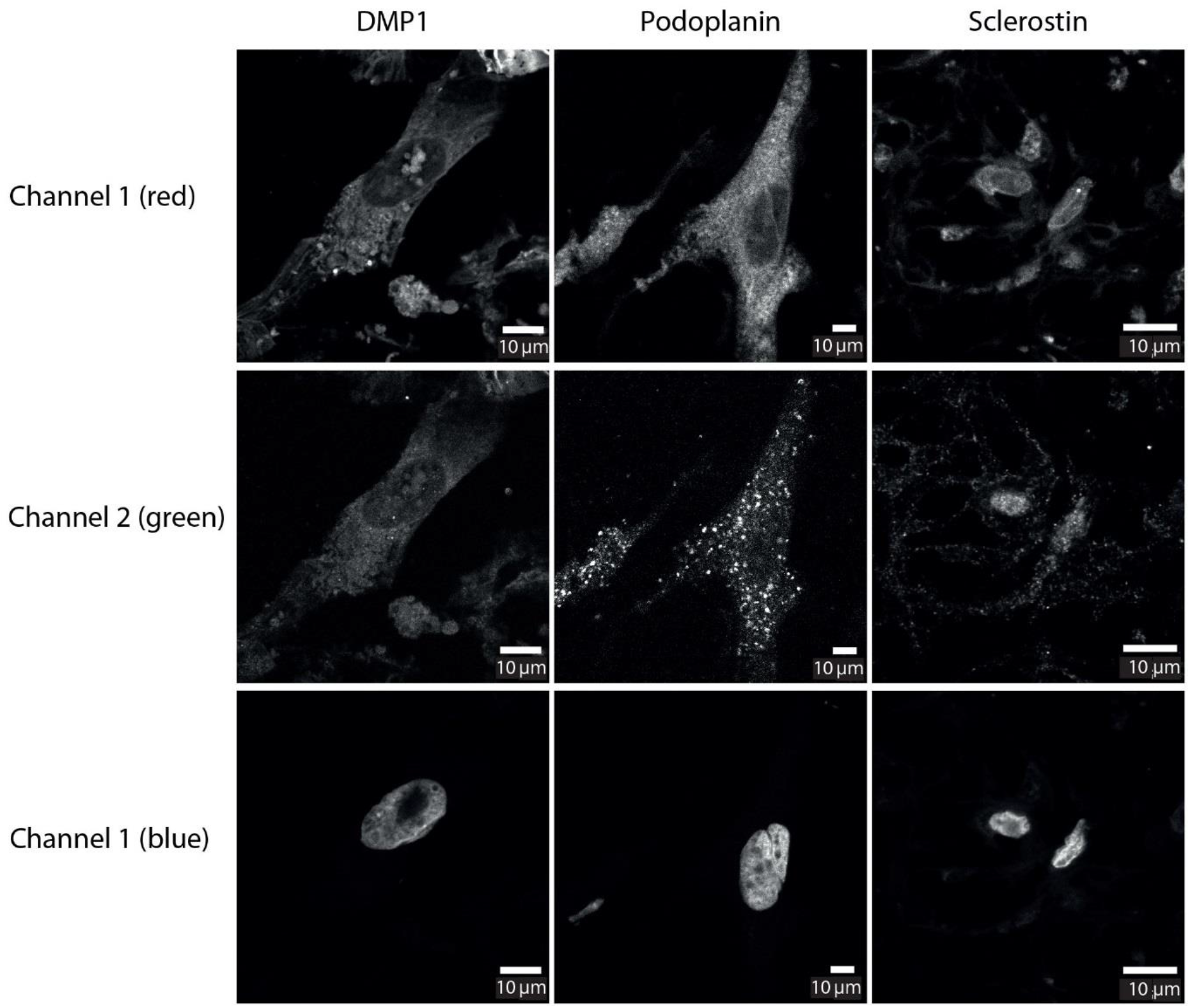
Separate channels of all panels from figure 1. The information of the three individual channels (Channel 1 - red, cell cytoplasm, Channel 2 - green, targeted protein and Channel 3 - blue, nuclei) are shown separately in gray scale for each image in Figure 1 g-i.

**Figure S5.**
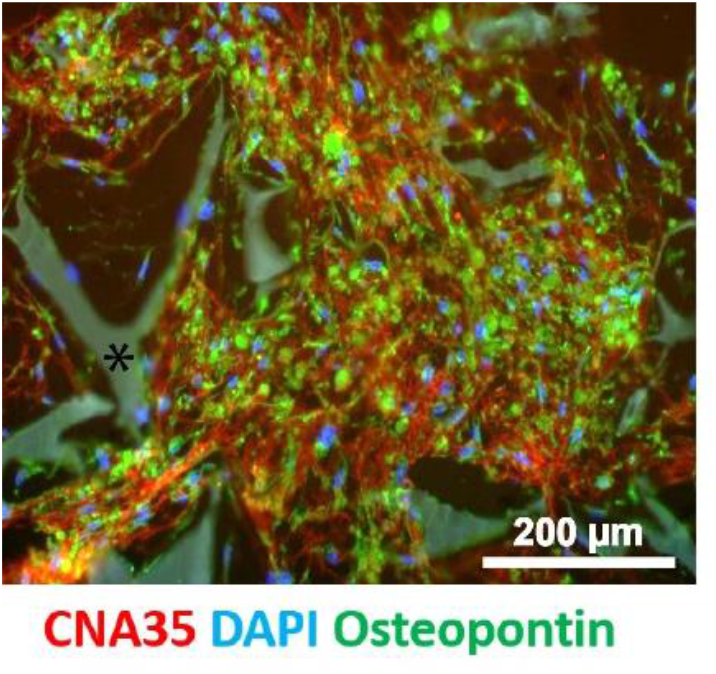
Co-expression of osteopontin and sclerostin. The image was taken from the same 3D co-culture sample as presented in figures 1k&l (after 8 weeks, see main text), at a different location within the scaffold. Osteopontin (green) production is indicated by its presence both in close proximity to the nuclei (DAPI, blue) and in the collagen matrix (red), implying the co-existence of osteoblasts and osteocytes organized in different domains.

**Figure S6.**
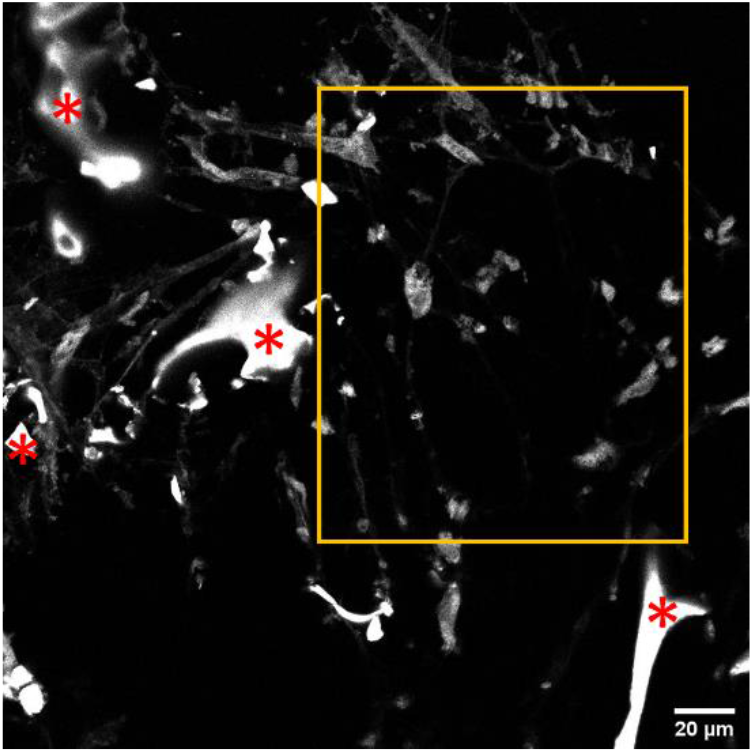
The original red channel image of Figure 2a without gamma correction. The cell cytoplasm is stained with FM 4-64. The image shows cells and silk fibroin (*) without gamma correction. The orange box indicates the area that is presented in Figure 2a with gamma correction.

**Figure S7.**
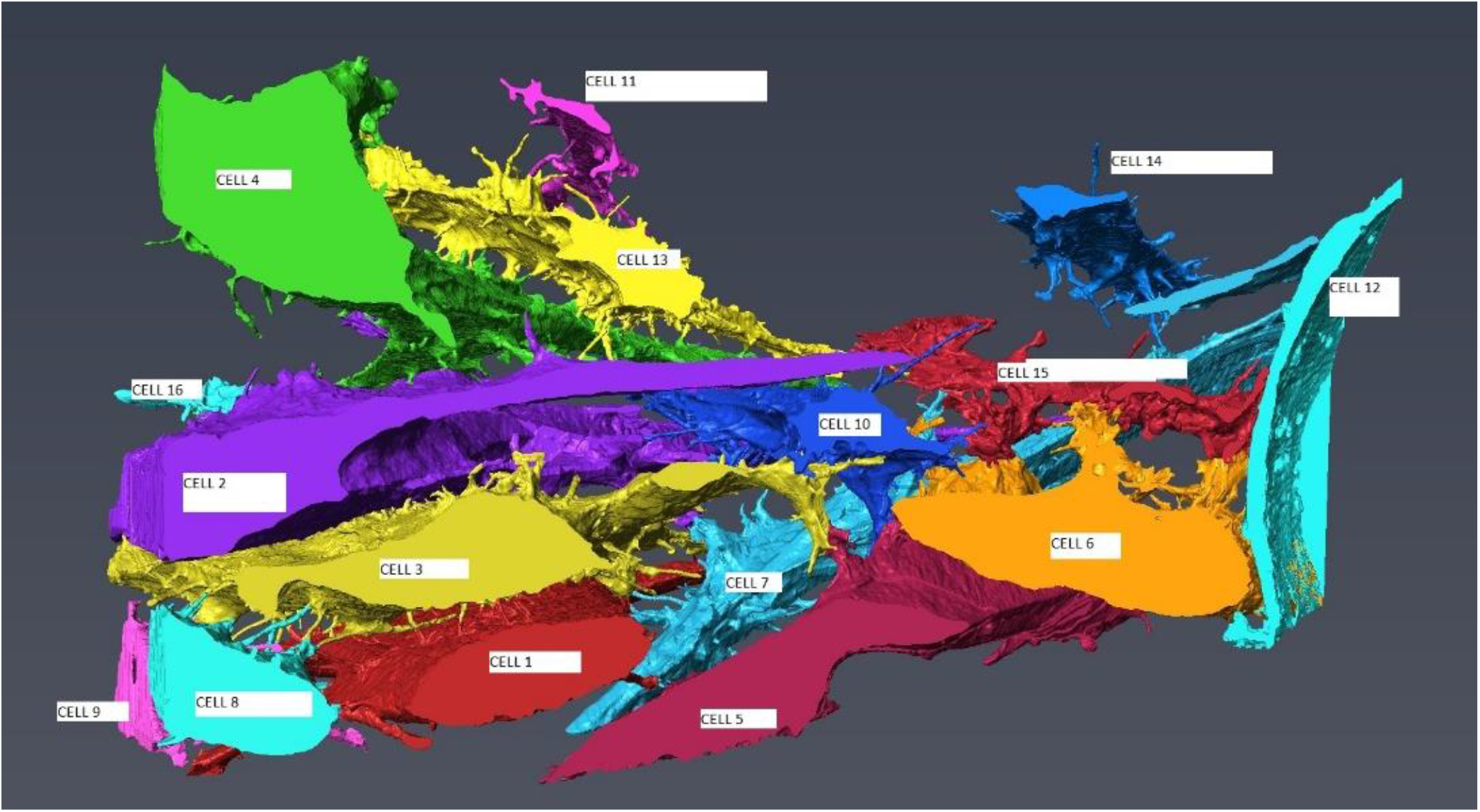
Cell number key map showing the different cells identified with the numbers corresponding to the numbers in Figure 2 and Tables S1 and S2.

**Figure S8.**
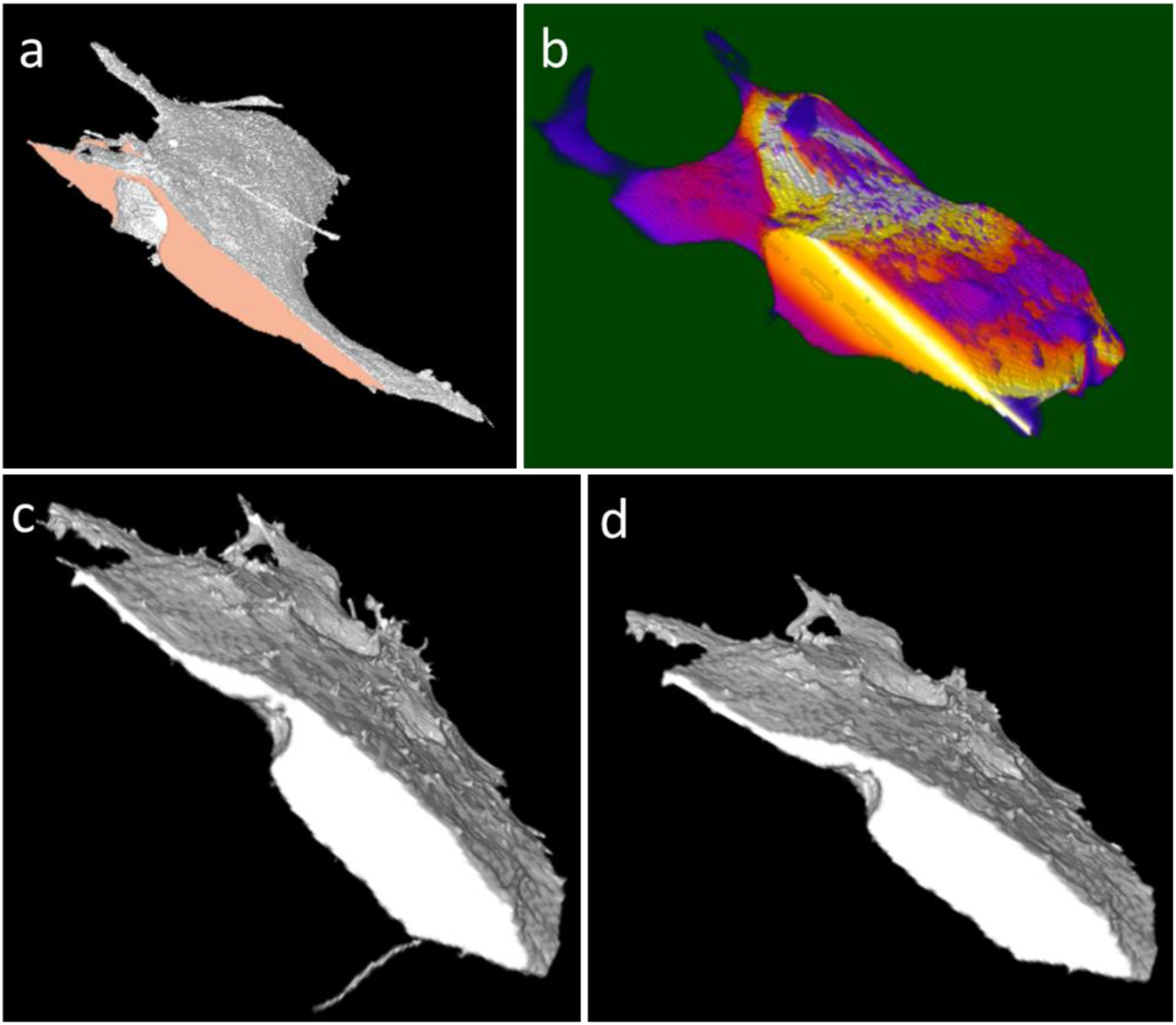
Image processing of cell surfaces. a) 3D rendering of a cell mask. The red area is the surface that is removed from the total area calculations as it is an artificial surface created by the edge of the image. b) The thickness measure of a cell mask as calculated by the plugin (see plugin desciption below). The dark purple areas depict a very small thickness measure shifting via red and orange to yellow and white for very large thickness areas. c-d) before and after view of the process removal plugin. c) view of a cell mask before applying the plugin, with some clear processes visible. d) the same cell mask after applying the plugin. The processes are no longer present.

### An Organoid for Woven Bone

For the image analysis we use the fact that processes are thin protrusions in order to detect and remove them. One can calculate the thickness of an image segment; however, using only the 3D thickness measure may create false positives as it gives the thinnest dimension at each point. A thin sheet of material can therefore not be distinguished from a cell process by this feature alone. Therefore the thickness of the segment in the x-, y-, and z-dimension are determined separately. Using the middle value of the thickness measures at each point we can detect areas that are thin in two dimensions without any restrictions on the third dimension. This separates the thin sheets of material from thin protrusions.

To calculate the cell process-to-cell surface unit ratio, several plugins in Fiji^[35]^ were applied to masks of the segmented cells created in Avizo using manual segmentation.

To calculate the surface area of each cell, the following sequence was use in Fiji:

1. ***Preprocessing***: The mask image was prepared for further processing by filling any internal holes, as they do not influence the surface area, but can create artifacts for the thickness measurements. To obtain a valid 3D thickness measure, the image was rescaled to equalize the x-, y-, and z-size of the voxels, creating an isotropic voxel size.
2. ***Determining observed cell surface***. (figure S8a): The total surface area of the cell mask is measured using the MorphoLibJ plugin.^[36]^ This plugin calculates the surface area computed using a discretized version of the Crofton formula, that computes intersections with line grids of thirteen orientations. Surface created by the edge of the image is detected and subtracted from the total surface area measurement.
3. ***3D thickness map***. (figure S8b): To calculate the thickness at all points in all dimensions, a separate thickness measurement was performed on each slice. After processing the whole stack, this procedure was repeated two times more after rotating the masked image subsequently around its x-axis and its y-axis.^[37]^ After combining the three thickness measures for each point, the maximum measured thickness was used as end value.
4. ***Removing processes***. (figure S8c-d): The resulting thickness image is now turned into a mask by setting a minimum threshold on the calculated thickness. Processes are removed by removing the areas with a thickness below this minimum.
5. ***Final surface area*** The surface area of the thickness mask is measured and any image edge surface is removed as in steps 3 and 4 above.

The plugin can be downloaded here: https://filesender.surf.nl/?s=download&token=e5f72e20-2240-4f58-abeb-3ecbeaab54f2

**Figure S9.**
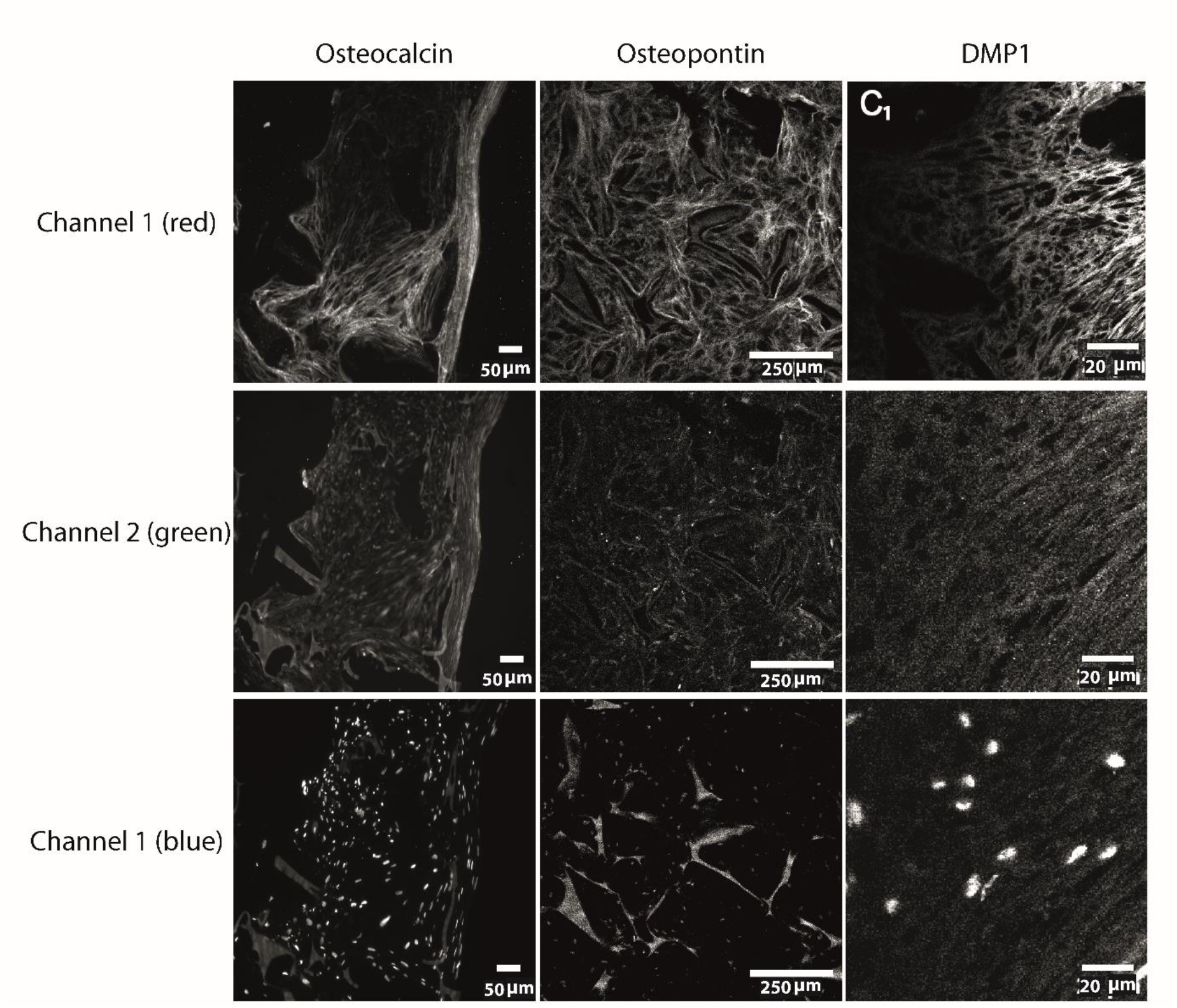
Separate channels of panels c-e from Figure 3. The information of the three individual channels (Channel 1 - red, cell cytoplasm, Channel 2 - green, targeted protein and Channel 3 -blue, nuclei) are shown in gray scale for each image in Figure 3.

**Figure S10.**
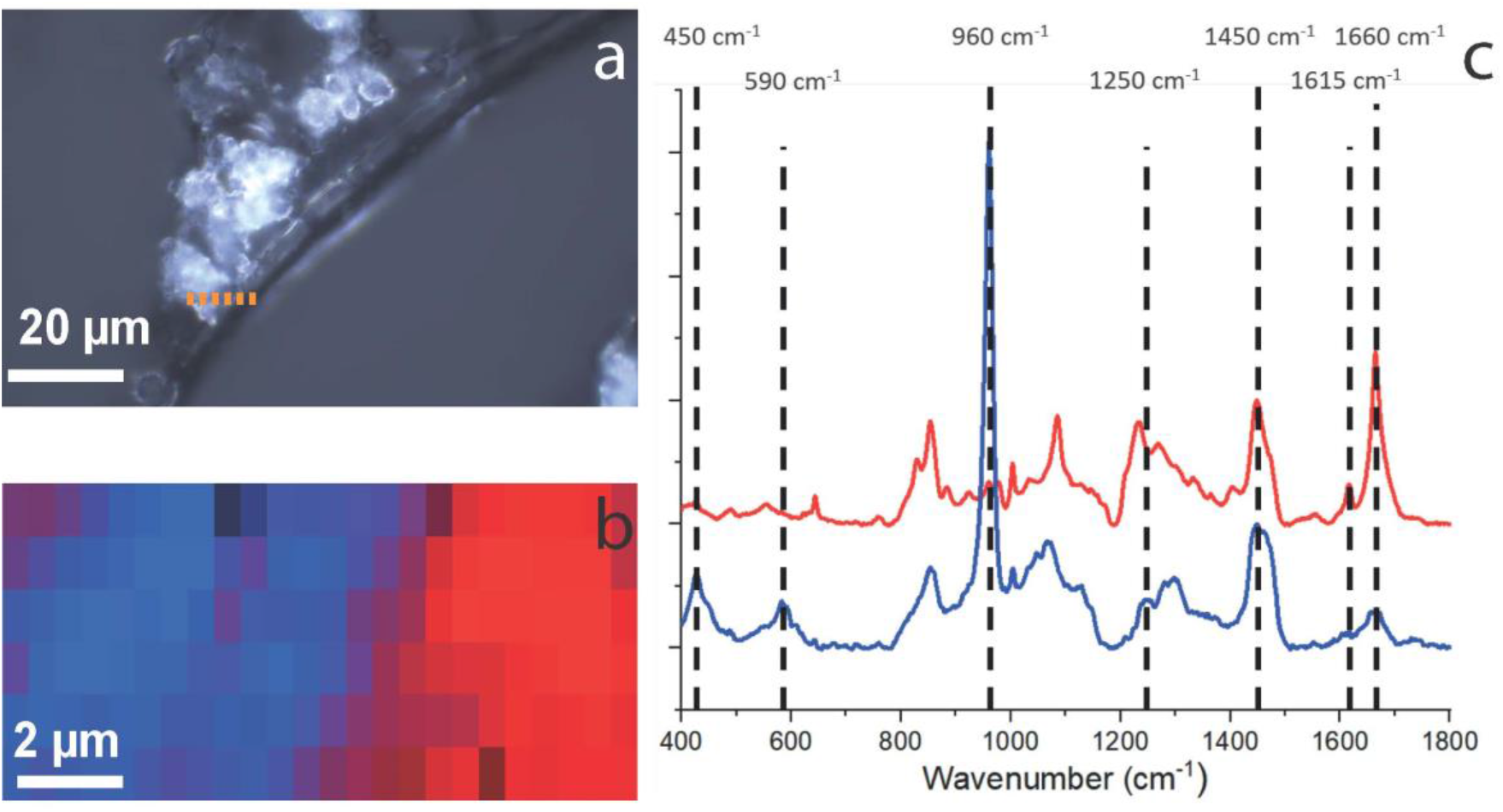
Raman micro-spectroscopy of the mineralized collagen - silk fibroin interface. a) Transmitted light image of the sample, the sample was sectioned into 10 micrometer thick layer and scanned using Raman microspectroscopy. The dotted line indicates the silk fibroin – mineralized collagen interface from which the depth scan of figure b was taken. b) Raman scan identifying different regions based on spectral differences, with the mineralized tissue in blue and the silk in red, the scanned area is 24 x 6 pixels. c) Corresponding spectra from b) (the Averaged spectra of the blue area shows colocalization of signals attributable to the mineral (450 cm^-1^: ν_2-_ PO_4_ vibration, 590 cm^-1^: ν_4-_ PO_4_ vibration, 960 cm^-1^ : ν_1-_ PO_4_ vibration) and the organic matrix (1250 cm^-1^: Amide III vibration; 1450 cm^-1^: CH_2_ bend vibration; 1660 cm^-1^: Amide I vibration). The red area is non-mineralized showing signals corresponding to the silk scaffold (1615 cm^-1^: Phenylalanine ring vibration; 1660 cm^-1^: Amide I vibration). The degree of mineralization - determined by taking the ratio of the phosphate ν_4_ (590 cm^-1^) intensity and the amide III (1250 cm^-1^) intensity – varied between 0.453-1.42 (average 1.17) in agreement with wat was found for woven bone (see figure 3g). After acquisition the Raman spectra were corrected for cosmic rays and background using the Project V plus software (WITec, Ulm). Background correction was performed using the shape fit option in the software, which corrects for the background based on the arc under the graph. The graphs were normalized to the 1450 cm^-1^ peak height. The Raman images were analyzed using the component analysis tool in the Project V plus software (WITec, Ulm). The spectral map was analyzed with two components, adding more components did not yield more chemically different areas, but could be used to indicate different mineralization levels within the measured area.

**Figure S11.**
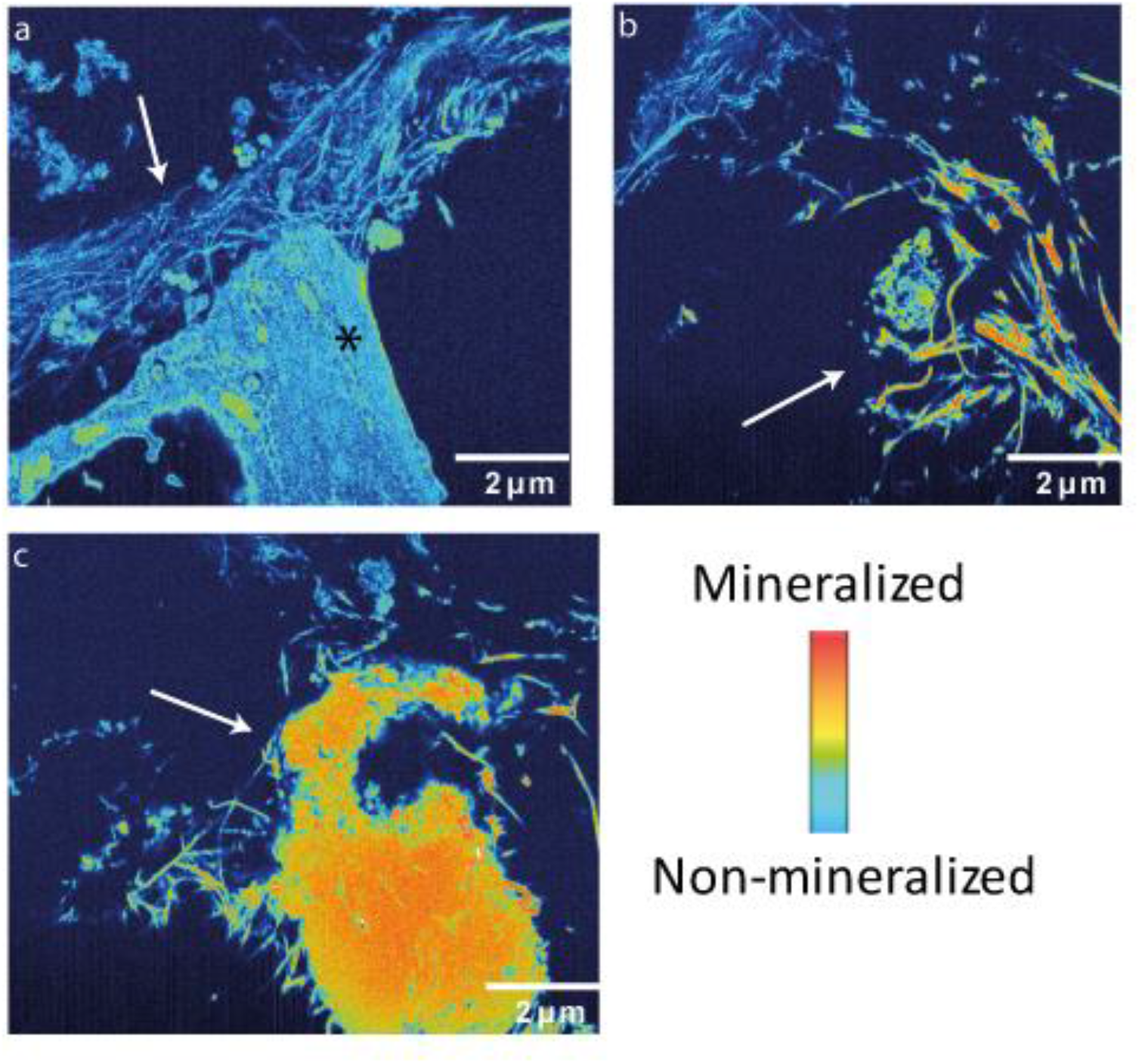
Development of the mineralized matrix: Z-slices from a 3D FIB/SEM co-culture sample after 5 weeks of culture, with heat map indicating the different levels of collagen mineralization a) unmineralized collagen fibrils (arrow) next to a cell (*) (slice #98, video S2). b) 3.24 µm deeper to the volume mineralized collagen is detected (arrow) (slice #260 video S2, 3.24µm. c) half micron deeper, the mineralized collagen fibrils evolve to form a large mineral precipitation (arrow) (slice #286, video S2). FIB/SEM Voxel size: 10×10×20 nm. The heat map is set with the collagen fibrils of lowest intensity (light blue, panel a) defined as non-mineralized collagen and the most dense structure (orange, panel c) defined as pure mineral precipitate.

**Table S1.** Number of processes per cell counted manually after segmentation of the 3D FIB/SEM volume in Video S1. The cell numbers correspond to the numbers shown in Figure S7.

|  | # of cell-cell<br>connecting<br>processes per<br>cell | # of<br>unconnected<br>processes per<br>cell | Total # of<br>processes per<br>cell |
| --- | --- | --- | --- |
| Cell 1 | 35 | 24 | 59 |
| Cell 2 | 46 | 24 | 70 |
| Cell 3 | 81 | 39 | 120 |
| Cell 4 | 23 | 23 | 46 |
| Cell 5 | 18 | 15 | 33 |
| Cell 6 | 7 | 11 | 18 |
| Cell 7 | 14 | 30 | 44 |
| Cell 8 | 21 | 3 | 24 |
| Cell 9 | 12 | 3 | 15 |
| Cell 10 | 15 | 27 | 42 |
| Cell 11 | 5 | 10 | 15 |
| Cell 12 | 4 | 15 | 19 |
| Cell 13 | 21 | 43 | 64 |
| Cell 14 | 0 | 40 | 40 |
| Cell 15 | 9 | 17 | 26 |

**Table S2.** Number of connections between cells counted in the volume manually after segmentation of the 3D FIB/SEM volume in Video S1. C = cell.

|  | C.<br>1 | C.<br>2 | C.<br>3 | C.<br>4 | C.<br>5 | C.<br>6 | C.<br>7 | C.<br>8 | C.<br>9 | C.<br>10 | C.<br>11 | C.<br>12 | C.<br>13 | C.<br>14 | C.<br>15 | Tot |
| --- | --- | --- | --- | --- | --- | --- | --- | --- | --- | --- | --- | --- | --- | --- | --- | --- |
| C. 1 | 0 |  | 26 |  |  |  | 4 | 3 | 2 |  |  |  |  |  |  | <u>35</u> |
| C. 2 |  | 0 | 39 | 2 |  |  | 2 |  |  | 2 |  | 1 |  |  |  | <u>46</u> |
| C. 3 | 26 | 39 | 0 |  | 1 |  | 1 | 7 | 2 | 5 |  |  |  |  |  | <u>81</u> |
| C. 4 |  | 2 |  | 0 |  |  |  |  |  | 5 |  |  | 16 |  |  | <u>23</u> |
| C. 5 |  |  | 1 |  | 0 | 6 | 6 | 4 |  | 1 |  |  |  |  |  | <u>18</u> |
| C. 6 |  |  |  |  | 6 | 0 |  |  |  | 2 |  |  |  |  | 5 | <u>13</u> |
| C. 7 | 4 | 2 | 1 |  | 6 |  | 0 |  |  |  |  |  |  |  | 1 | <u>14</u> |
| C. 8 | 3 |  | 7 |  | 4 |  |  | 0 | 8 |  |  |  |  |  |  | <u>22</u> |
| C. 9 | 2 |  | 2 |  |  |  |  | 8 | 0 |  |  |  |  |  |  | <u>12</u> |
| C. 10 |  | 2 | 5 | 5 | 1 | 2 |  |  |  | 0 |  |  |  |  |  | <u>15</u> |
| C. 11 |  |  |  |  |  |  |  |  |  |  | 0 |  | 5 |  |  | <u>5</u> |
| C. 12 |  | 1 |  |  |  |  |  |  |  |  |  | 0 |  |  | 3 | <u>4</u> |
| C. 13 |  |  |  | 16 |  |  |  |  |  |  | 5 |  | 0 |  |  | <u>21</u> |
| C. 14 |  |  |  |  |  |  |  |  |  |  |  |  |  | 0 |  | <u>0</u> |
| C. 15 |  |  |  |  |  | 5 | 1 |  |  |  |  | 3 |  |  | 0 | <u>9</u> |
| **Tot** | <u>35</u> | <u>46</u> | <u>81</u> | <u>23</u> | <u>18</u> | <u>13</u> | <u>14</u> | <u>22</u> | <u>12</u> | <u>15</u> | <u>5</u> | <u>4</u> | <u>21</u> | <u>0</u> | <u>9</u> |  |

**Table S3.** Calculation of number of processes per cell surface unit. Numbers of processes taken from table S1 (Total # of processes per cell), processes were counted manually. Cell surface area was calculated as described in figure S8.

| Cell No. | (n) # total processes | (S) surface area [$\mu\text{m}^2$] | n/S |
| --- | --- | --- | --- |
| cell 1 | 59 | 728 | 0,081 |
| cell 2 | 70 | 1370 | 0,051 |
| cell 3 | 120 | 989 | 0,121 |
| cell 4 | 46 | 663 | 0,069 |
| cell 5 | 33 | 628 | 0,052 |
| cell 6 | 24 | 341 | 0,070 |
| cell 7 | 44 | 791 | 0,055 |
| cell 8 | 25 | 149 | 0,167 |
| cell 9 | 15 | 224 | 0,066 |
| cell 10 | 42 | 244 | 0,172 |
| cell 11 | 15 | 92 | 0,162 |
| cell 12 | 19 | 253 | 0,075 |
| cell 13 | 64 | 599 | 0,106 |
| cell 14 | 40 | 171 | 0,234 |
| cell 15 | 26 | 383 | 0,067 |

### Video S1

Video S1 shows a three dimensional reconstruction and segmentation of the 3D FIB/SEM stack imaging an area of 40 µm x 20 µm x 20 µm of the 3D co-culture after 5 weeks of culture. This movie shows 15 cells that all show at least 15 processes, of which some are connected to other cells (see Figure 3, Figure S6 and Table S1 and S2 for more information). The cells are embedded in a dense collagen matrix (cyan). The stack is composed of 316 images, taken with a voxel size of 30 nm x 30 nm x 30 nm. A high resolution version of the movie is available upon request

### Video S2

**Video S2**. Development of the mineralized matrix in 3D. 3D FIB-SEM stack taken with a voxel size of 10×10×20 nm, showing a volume of 16×13×6 micrometer^3^. The sample starts with showing a large uncontrolled mineral precipitate (orange). A 3 micrometer layer of non-mineralized collagen appears at the upper part of the movie (light blue fibrils). The non-mineralized collagen located in close proximity to a cell. When the mineral precipitate fades, collagen fibrils with strong orange-red color are revealed at the right side. These fibrils evolve to form a large mineralized structure. This could indicate mineral deposition that started under biological control, by mineralizing single collagen fibrils, but growing to form a larger mineral precipitate. For more information see Fig. 3h and Figure S11. A high resolution version of the movie is available upon request.

## References

[1] Principles of Bone Biology (Fourth Edition), Eds John P. Bilezikian, T. John Martin, Thomas L. Clemens, & Clifford J. Rosen, 1943-1986 (Academic Press, 2020).

[2] A. Fatehullah, S. H. Tan, N. Barker, Nat Cell Biol 2016, 18, 246–254.

[3] L. F. Bonewald, J Bone Miner Res 2011, 26, 229–238.

[4] T. A. Franz-Odendaal, B. K. Hall, P. E. Witten, Dev Dynam 2006, 235, 176–190.

[5] C. Zhang, A. D. Bakker, J. Klein-Nulend, N. Bravenboer, Curr Osteoporos Rep 2019, 17, 207–216.

[6] aK. Wang, L. Le, B. M. Chun, L. M. Tiede-Lewis, L. A. Shiflett, M. Prideaux, R. S. Campos, P. A. Veno, Y. Xie, V. Dusevich, L. F. Bonewald, S. L. Dallas, J Bone Miner Re 2019, 34, 979-995; bA. Iordachescu, H.D. Amin, S.M. Rankin, R.L. Williams, C. Yapp, A. Bannerman, A. Pacureanu, O. Addison, P.A. Hulley, L.M. Grover, Advanced Biosystems 2018, 2.

[7] aG. Thrivikraman, A. Athirasala, R. Gordon, L. Zhang, R. Bergan, D. R. Keene, J. M. Jones, H. Xie, Z. Chen, J. Tao, B. Wingender, L. Gower, J. L. Ferracane, L. E. Bertassoni, Nat Commun 2019, 10, 3520; bG. Nasello, P. Alamán-Díez, J. Schiavi, M.Á. Pérez, L. McNamara, J. M. García-Aznar, Frontiers in Bioengineering and Biotechnology 2020, 8.

[8] J. Skottke, M. Gelinsky, A. Bernhardt, Int J Mol Sci 2019, 20.

[9] J. R. Vetsch, S. J. Paulsen, R. Muller, S. Hofmann, Acta Biomater 2015, 13, 277–285.

[10] M. Kerschnitzki, A. Akiva, A. Ben Shoham, Y. Asscher, W. Wagermaier, P. Fratzl, L. Addadi, S. Weiner, J Struct Biol 2016, 195, 82–92.

[11] J. Melke, F. Zhao, B. van Rietbergen, K. Ito, S. Hofmann, Eur Cells Mater 2018, 36, 57–68.

[12] R.L.v.B. Kenneth E.S. Poole, Nigel Loveridge, Herman Hamersma,, C.W.L. Socrates E. Papapoulos, and Jonathan Reeve, The FASEB Journal 2005, 19, 1842–1844.

[13] S. W. Verbruggen, T. J. Vaughan, L. M. McNamara, J R Soc Interface 2012, 9, 2735–2744.

[14] P. Varga, B. Hesse, M. Langer, S. Schrof, N. Mannicke, H. Suhonen, A. Pacureanu, D. Pahr, F. Peyrin, K. Raum, Biomech Model Mechanobiol 2015, 14, 267–282.

[15] M. Kerschnitzki, W. Wagermaier, P. Roschger, J. Seto, R. Shahar, G. N. Duda, S. Mundlos, P. Fratzl, J Struct Biol 2011, 173, 303–311.

[16] R. Yair, A. Cahaner, Z. Uni, R. Shahar, Poult Sci 2017, 96, 2301–2311.

[17] D. Tokarz, R. Cisek, M. N. Wein, R. Turcotte, C. Haase, S. A. Yeh, S. Bharadwaj, A. P. Raphael, H. Paudel, C. Alt, T. M. Liu, H. M. Kronenberg, C. P. Lin, PLoS One 2017, 12, e0186846.

[18] S. W. and, H. D. Wagner, Annual Review of Materials Science 1998, 28, 271–298.

[19] N. Reznikov, R. Shahar, S. Weiner, Bone 2014, 59, 93–104.

[20] N. Reznikov, R. Shahar, S. Weiner, Acta Biomaterialia 2014, 10, 3815–3826.

[21] B. W. M. de Wildt, S. Ansari, N. A. J. M. Sommerdijk, K. Ito, A. Akiva, S. Hofmann, Current Opinion in Biomedical Engineering 2019, 10, 107–115.

[22] M. Kazanci, P. Roschger, E. P. Paschalis, K. Klaushofer, P. Fratzl, Journal of Structural Biology 2006, 156, 489–496.

[23] A. Akiva, M. Kerschnitzki, I. Pinkas, W. Wagermaier, K. Yaniv, P. Fratzl, L. Addadi, S. Weiner, J Am Chem Soc 2016, 138, 14481–14487.

[24] M. D. McKee, A. Nanci, Microscopy Research and Technique 1995, 31, 44–62.

[25] S. Weiner, W. Traub, FEBS letters 1986, 206, 262–266.

[26] S. E. Park, A. Georgescu, D. Huh, Science 2019, 364, 960–965.

[27] S. Hofmann, H. Hagenmuller, A. M. Koch, R. Muller, G. Vunjak-Novakovic, D. L. Kaplan, H. P. Merkle, L. Meinel, Biomaterials 2007, 28, 1152–1162.

[28] R. Nazarov, H.-J. Jin, D. L. Kaplan, Biomacromolecules 2004, 5, 718–726.

[29] aL. A. Hildebrand T, Muller R, Dequeker J, Ruegsegger P., J. Bone Miner. Res. 1999, 14; bG. H. van Lenthe, H. Hagenmuller,M. Bohner, S.J. Hollister, L. Meinel, R. Muller, Biomaterials 2007, 28, 2479–2490.

[30] J. Fermie, N. Liv, C. Ten Brink, E. G. van Donselaar, W. H. Muller, N. L. Schieber, Y. Schwab, H. C. Gerritsen, J. Klumperman, Traffic 2018, 19, 354–369.

[31] aS. T. Oorschot VMJ, Bryson-Richardson RJ, Ramm G., in Correl. Light Electron Microsc. II Academic Press, 2014, pp. 241-258; bE. van Meel, M. Boonen, H. Zhao, V. Oorschot, F. P. Ross, S. Kornfeld, J. Klumperman, Traffic 2011, 12, 912–924.

[32] J. Slot, Geuze, H., Nat protoc 2007, 2, 2480–2491.

[33] J. S. Nyman, A. J. Makowski, C. A. Patil, T. P. Masui, E. C. O’Quinn, X. Bi, S. A. Guelcher, D. P. Nicollela, A. Mahadevan-Jansen, Calcified Tissue International 2011, 89, 111–122.

[34] K. N. Krahn, C. V. C. Bouten, S. van Tuijl, M. A. M. J. van Zandvoort, M. Merkx, Analytical Biochemistry 2006, 350, 177–185.

[35] J. Schindelin, I. Arganda-Carreras, E. Frise, V. Kaynig, M. Longair, T. Pietzsch, S. Preibisch, C. Rueden, S. Saalfeld, B. Schmid, J.-Y. Tinevez, D. J. White, V. Hartenstein, K. Eliceiri, P. Tomancak, A. Cardona, Nature Methods 2012, 9, 676–682.

[36] D. Legland, I. Arganda-Carreras, P. Andrey, Bioinformatics 2016, 32, 3532–3534.

[37] E. H. W. Meijering, W. J. Niessen, M. A. Viergever, Medical Image Analysis 2001, 5, 111–126.

